## Supplementary Information for "Master corepressor inactivation through multivalent SLiM-induced polymerization mediated by the oncogene suppressor RAI2"

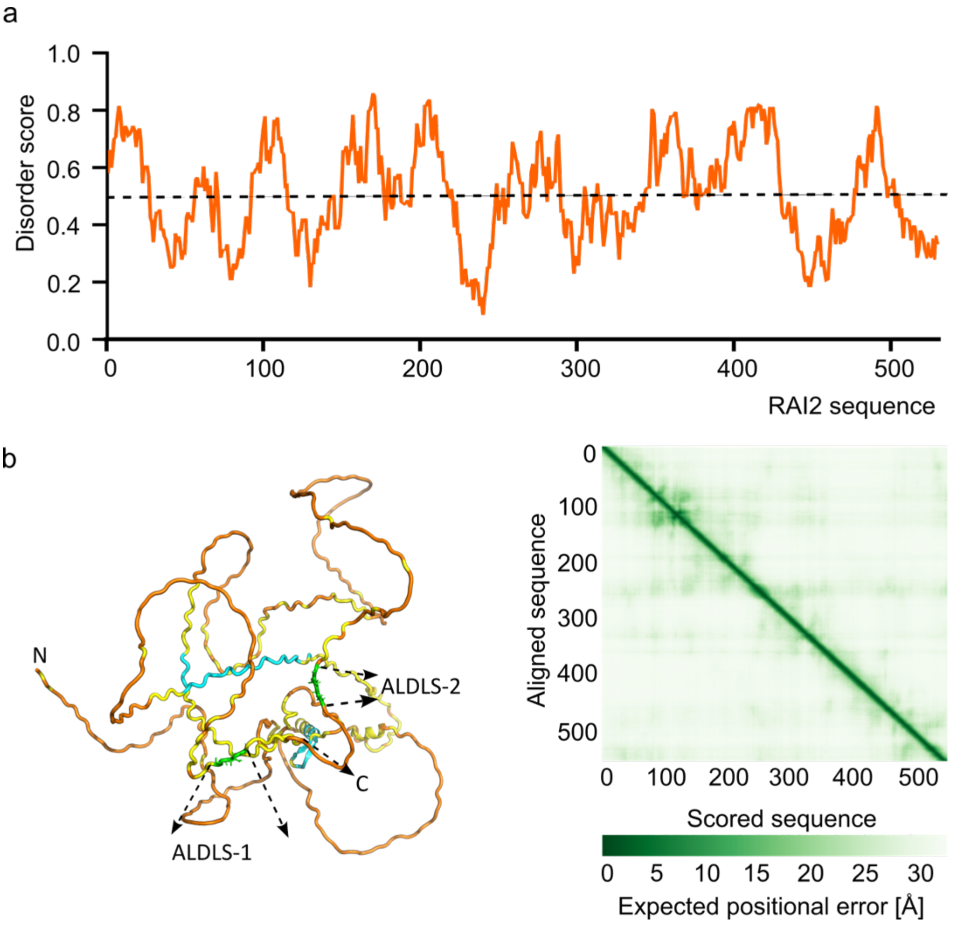

**Supplementary Fig. 1: Secondary and tertiary structure prediction of the RAI2 protein.** a. Prediction of disordered protein sequence regions by IUPRED disorder score in orange [1,2]. IUPRED score > 0.5 (indicated by dashed line) points towards intrinsically disordered sequence regions. b. Left, RAI2 structure prediction by AlphaFold [3-5] indicates a lack of tertiary structure for most of the RAI2 sequence. The RAI2 sequence is colored according to the per-residue confidence score (pLDDT): 90–70, cyan; 69–50, yellow; <50, orange. The two ALDLS motifs are in green and labeled. The pLDDT score for ALDLS-1 is between 50 and 70, while for ALDLS-2 it is less than 50, indicating low confidence in structure prediction. The N- and C-termini are indicated with “N” and “C”. Right, high distance errors [Å] indicated in the Predicted Aligned Error (PAE) plot confirm the absence of folded domains and the lack of tertiary structure.

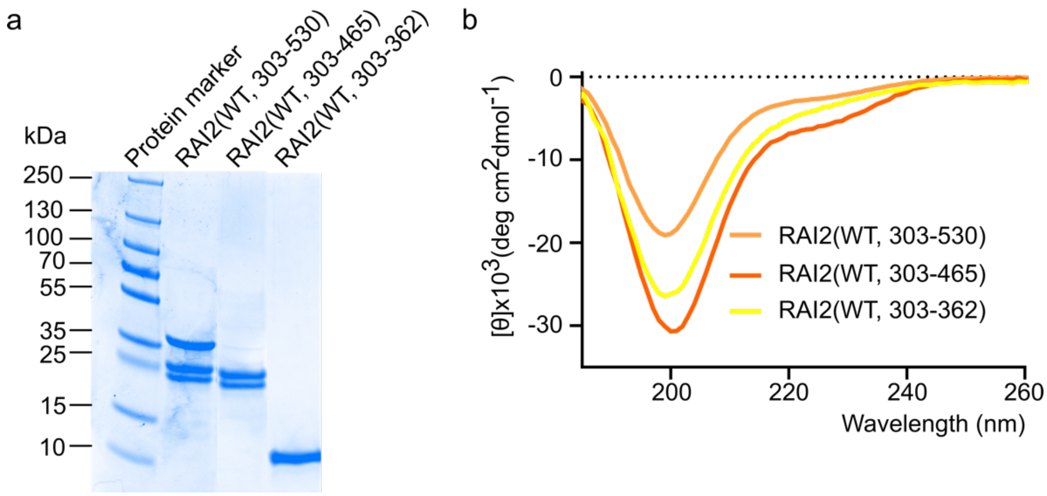

**Supplementary Fig. 2**: **Characterization of the truncated RAI2 protein constructs (*cf.* Fig.1a)**. a. SDS-PAGE analysis, molecular weight markers are indicated on the left. RAI2(WT, 303-465) and RAI2(WT, 303-530) show a significant tendency for degradation at the residue S467, as evidenced by mass spectrometry, whereas RAI2(WT, 303-362) is proteolytically stable in solution. The procedure and parameters used for mass spectrometry analysis have been described in the Supplementary Methods and the source data are provided in Supplementary Data 8. b. CD spectra of RAI2 truncated constructs used for biophysical and structural analysis. All CD spectra indicate the absence of a significant amount of secondary structure.

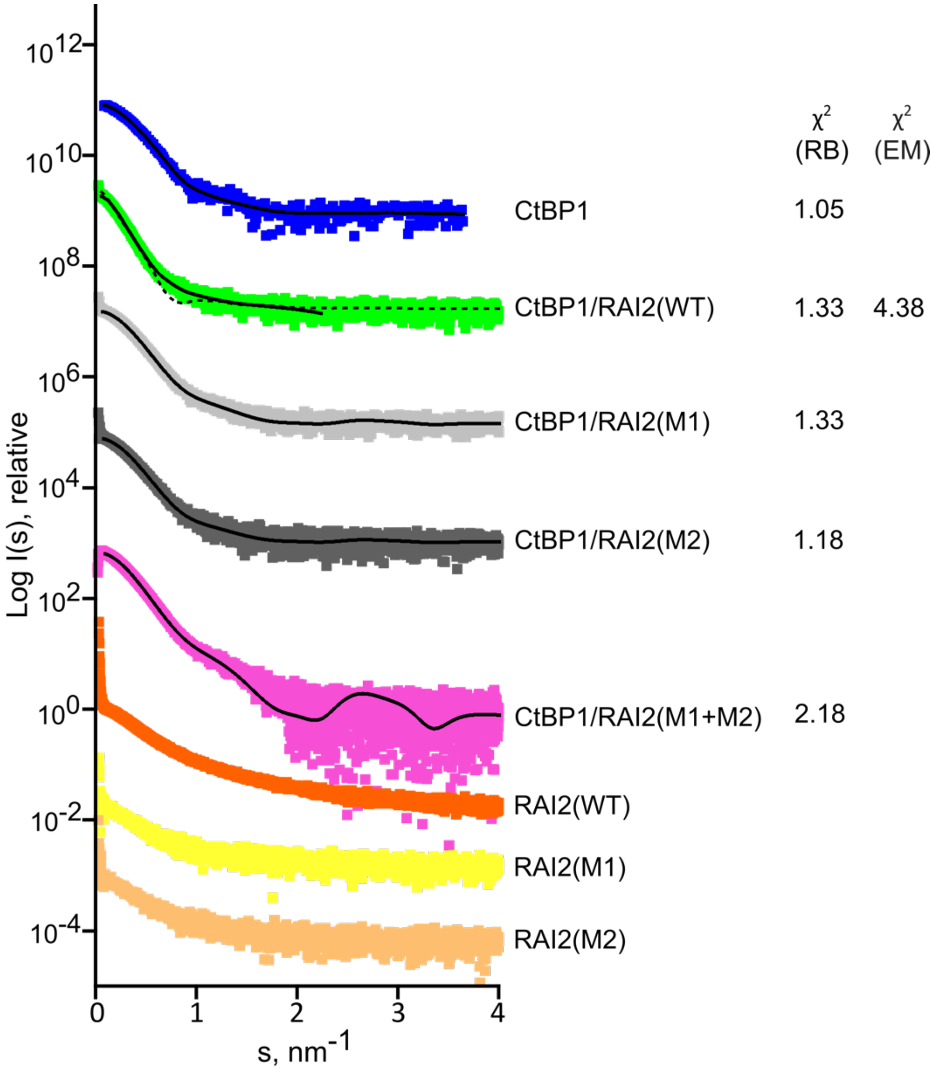

**Supplementary Fig. 3: SAXS data and model fits for CtBP1, RAI2(303-465) variants and CtBP1/RAI2(303-465) variant complexes (shown as squares).** Rigid body (RB, solid lines) and cryo-EM model (dashed lines) fits to the experimental SAXS data are shown. The corresponding χ2 statistics are listed on the right. The plots are based on a single experiment and the error bars represent the SEM on the scattering intensities measured at each point in s following Poisson counting statistics [6]. In some of the samples, an increase in scattering intensity at very low scattering angles has been observed. This feature is most likely due to aggregated fractions caused by rapid radiation damage of the sample [7]. The data have been taken from SASBDB entries SASDQW5, SASDQC6, SASDQD6, SASDQE6, SASDQF6, SASDQ46, SASDQ56 and SASDQ66. See also Supplementary Data 1 and 5.

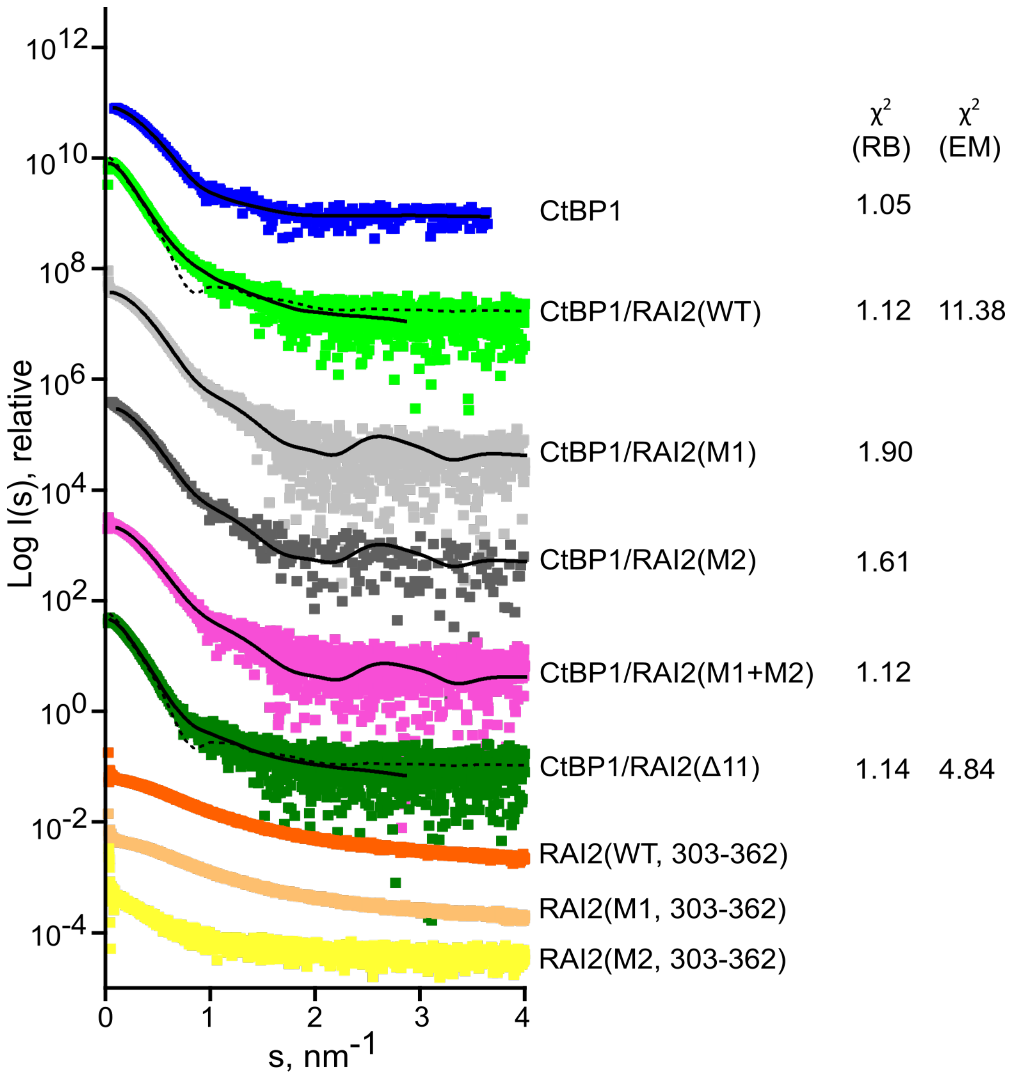

**Supplementary Fig. 4: SAXS data and model fits for CtBP1, RAI2(303-362) variants and CtBP1/RAI2(303-362) variant complexes (shown as squares).** Fits of rigid body(RB, solid black lines) and cryo-EM models (dashed lines) to the experimental SAXS data are shown. The χ2 goodness-of-fit associated with each fit is shown to the right of each data set. The plots are based on a single experiment and the error bars represent the SEM on the scattering intensities measured at each point in s following Poisson counting statistics [6]. In some of the samples, an increase in scattering intensity at very low scattering angles has been observed. This feature is most likely due to aggregated fractions caused by rapid radiation damage of the sample [7]. The data have been taken from SASBDB entries SASDQW5, SASDQ76, SASDQ86, SASDQ96, SASDQA6, SASDQB6, SASDQZ5, SASDQ26 and SASDQ36 forms the basis for the scattering plots. See also Supplementary Data 2 and 6.

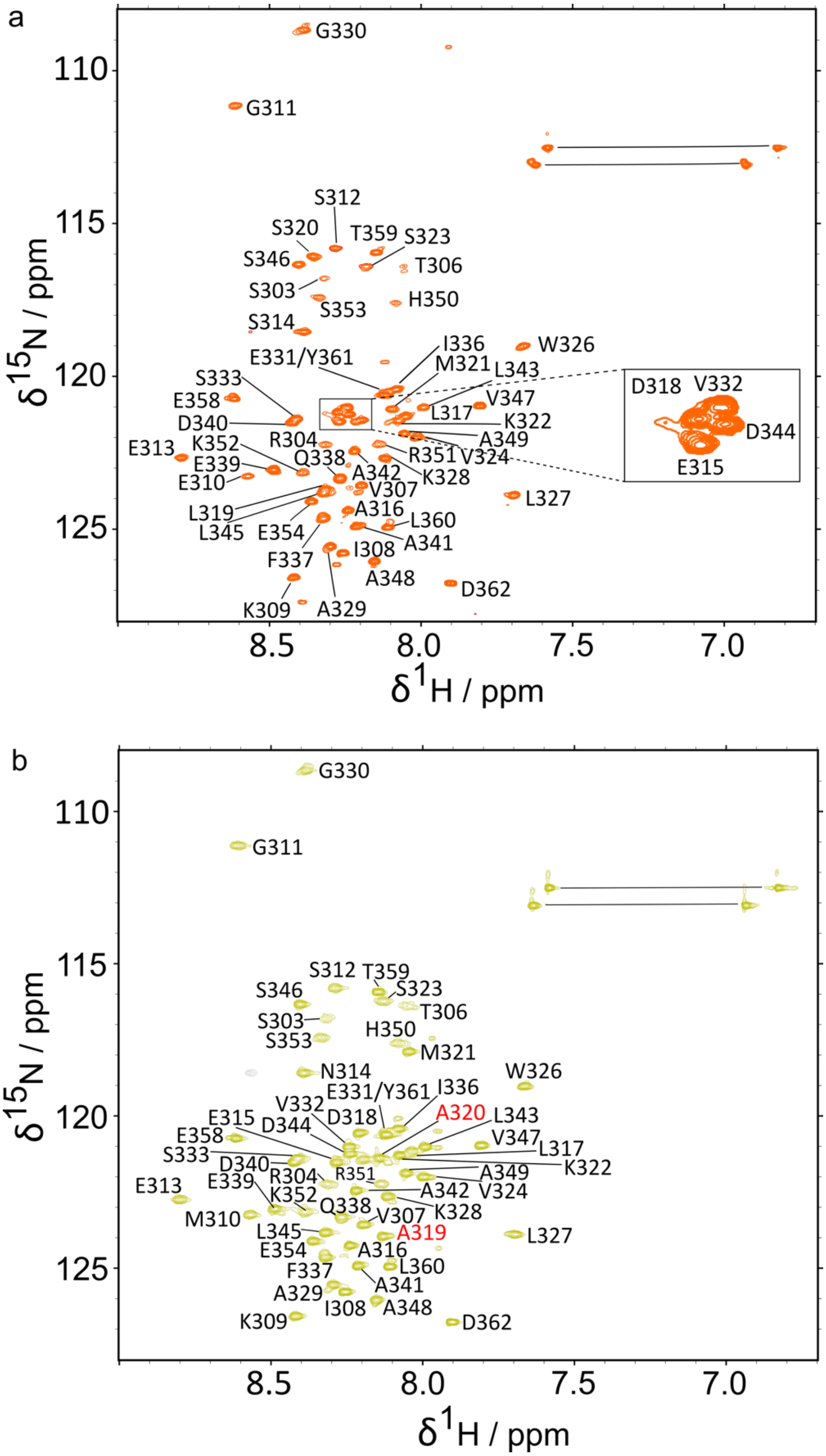
a
G330
G311
S312
S320
T359
S323
S346
T306
S303
S353
H350
S314
W326
E331/Y361
I336
M321
L343
S333
D318
V332
E358
V347
D340
L317
K322
D344
E315
A349
R304
E313
K352
R351
V324
Q338
E339
A342
K328
E310
V307
L327
L319
A316
L345
E354
L360
A341
I308
F337
A348
D362
K309
A329
110
115
120
125
8.5
8.0
7.5
7.0
δ15N / ppm
δ1H / ppm
b
G330
G311
S312
T359
S346
S323
T306
S303
H350
S353
M321
N314
W326
E331/Y361
I336
V332
A320
L343
E315
D344
D318
E358
V347
L317
S333
D340
K322
R304
R351
A349
E313
E339
K352
A342
V324
Q338
M310
V307
K328
L345
A319
E354
A316
F337
L360
A341
A329
I308
A348
K309
D362
110
115
120
125
8.5
8.0
7.5
7.0
δ15N / ppm
δ1H / ppm

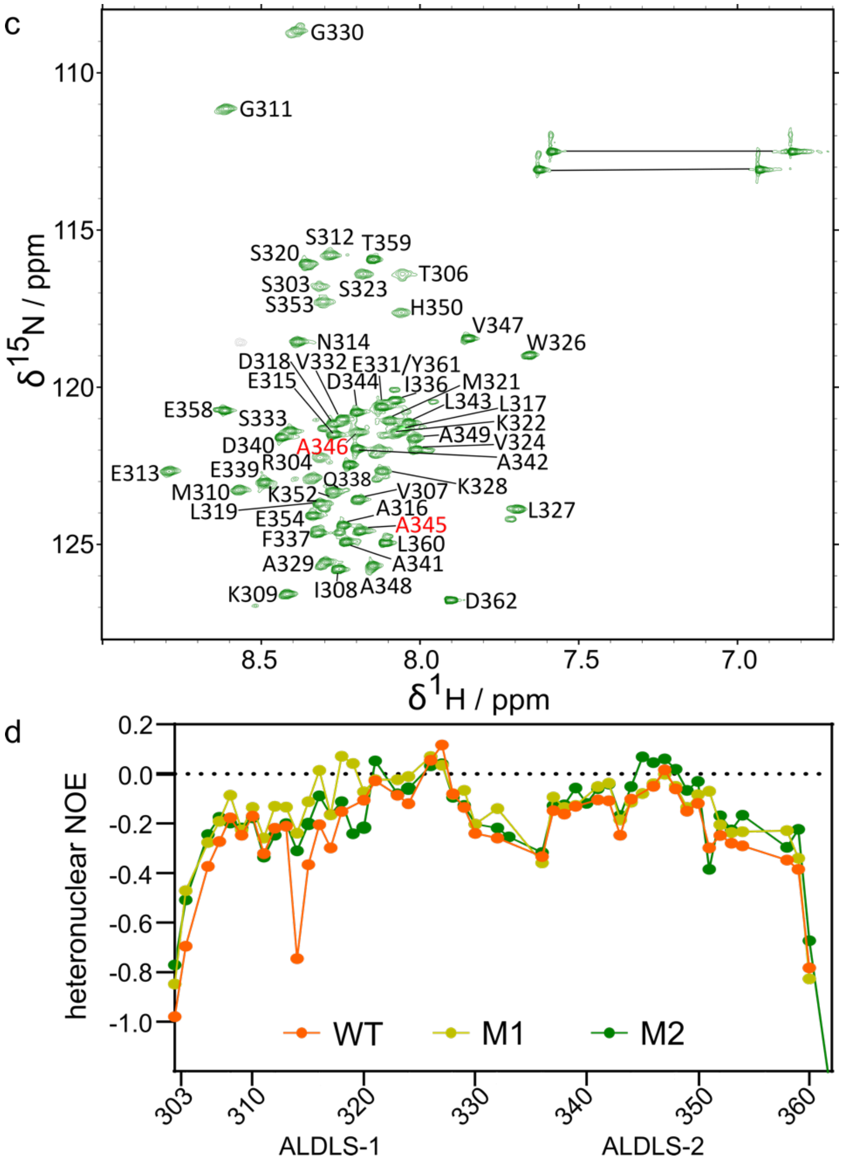

**Supplementary Fig. 5: NMR characterization of RAI2(303-362) variants.** [$^{1}$H,$^{15}$N]-HSQC spectra of the $^{15}$N-labeled RAI2(WT) in orange (a), RAI2(M1) in green-yellow (b) and RAI2(M2) in dark green (c). Reduced spectral peak dispersion within 1 ppm demonstrate that all RAI2 variants are intrinsically disordered. Assignments for the backbone amides are annotated and are listed in Supplementary Data 1. Mutated residues in RAI2 M1 and M2 variants are annotated in red. The most crowded part of the HSQC of RAI2(WT) is also shown enlarged in the inserted box for annotation purposes. Non-degenerate protons of side chain amino groups are connected by the straight lines. d. $^{15}$N-{$^{1}$H}-heteronuclear NOE ratio of RAI2(WT), RAI2(M1) and RAI2(M2), indicating fast time scale mobility of backbone HN bonds along the sequence. Negative heteronuclear NOE values confirm the disordered nature of RAI2 variants. The two RAI2 ALDLS motifs are annotated. Residue numbers of RAI2(303-362) are indicated. Source data are provided as a Source Data file.

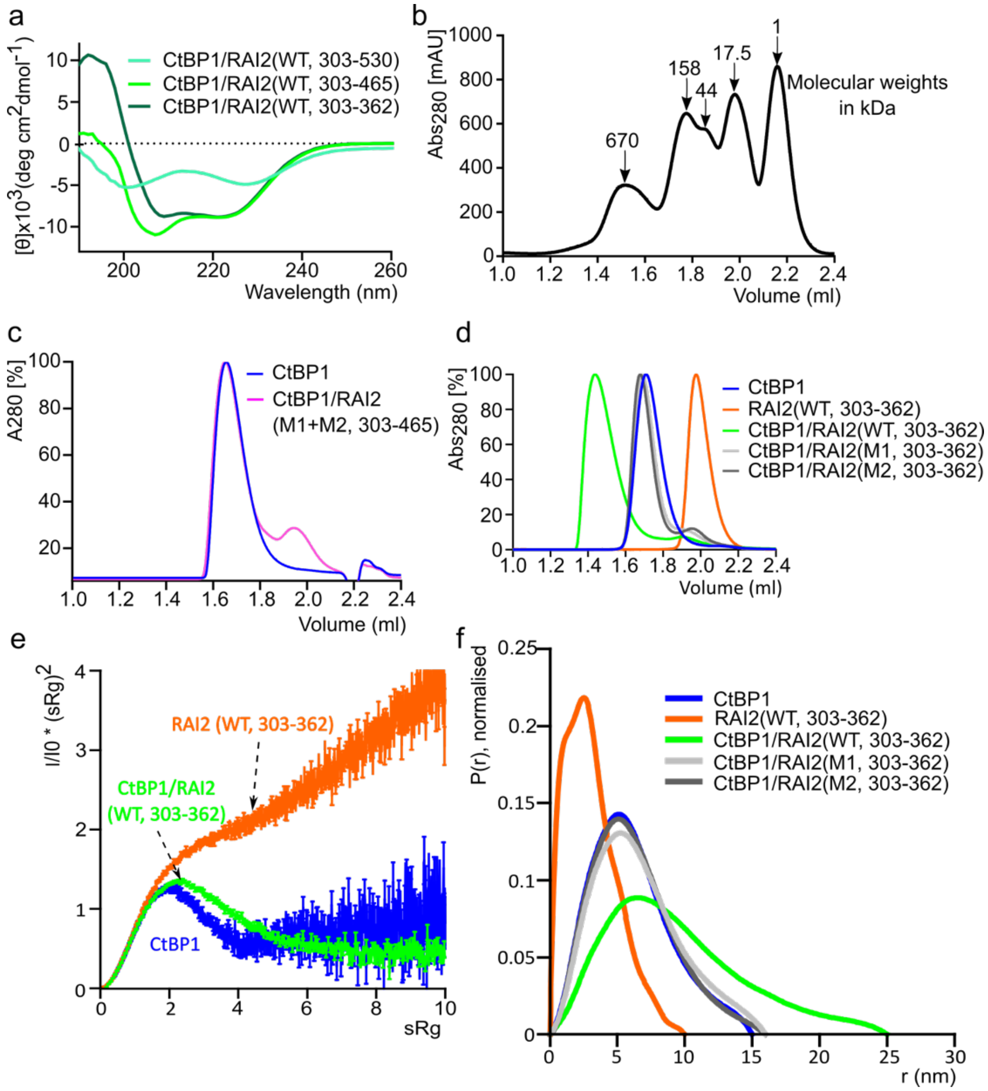

**Supplementary Fig. 6: Biophysical characterization of the CtBP1/RAI2(303-362) complexes.** a. CD spectra depicting the complexes of CtBP1 with RAI2 protein constructs of different lengths (*cf*. Fig.1a). The data shown for CtBP1/RAI2(WT, 303-465) are identical to those shown in Fig.2b. b. Superose 6 SEC profile of protein markers on a 2.4 ml column containing five standard proteins: thyroglobulin (670 kDa), gamma globulin (158 kDa), ovalbumin (44 kDa), myoglobin (17.5 kDa) and vitamin B12 (1.35 kDa). c. Normalized SEC profile after titration of RAI2(M1+M2) with CtBP1(WT) (magenta). CtBP1(WT) SEC profile is in blue. RAI2(M1+M2) shows no evidence of complex formation with CtBP. d. Normalized SEC profiles after titration of the RAI2(WT, green), RAI2(M1, light grey) and RAI2(M2, dark grey) with CtBP1. SEC profiles of separate RAI2(WT, orange) and CtBP1 (blue) are also shown. The measured retention volumes are listed in Supplementary Table 2. e. SAXS Kratky plot showing CtBP1(WT, blue), RAI2(WT, orange) and CtBP1(WT)/RAI2(WT) complex in green. The plots are based on a single experiment and the error bars represent the propagated SEM on the scattering intensities measured at each point in s following Poisson counting statistics [6]. The data shown for CtBP1(WT) are identical to those shown in Fig.1c. The data have been taken from SASBDB entries SASDQW5, SASDQZ5 and SASDQ76. f. SAXS distance distribution P(r) profiles of CtBP1(WT), RAI2(WT), and complexes of CtBP1 with RAI2(WT), RAI2(M1) and RAI2(M2), using color conventions used elsewhere. The plots are based on a single experiment and the data have been taken from SASBDB entries SASDQW5, SASDQZ5, SASDQ76, SASDQ86 and SASDQ96. Source data are provided as a Source Data file.

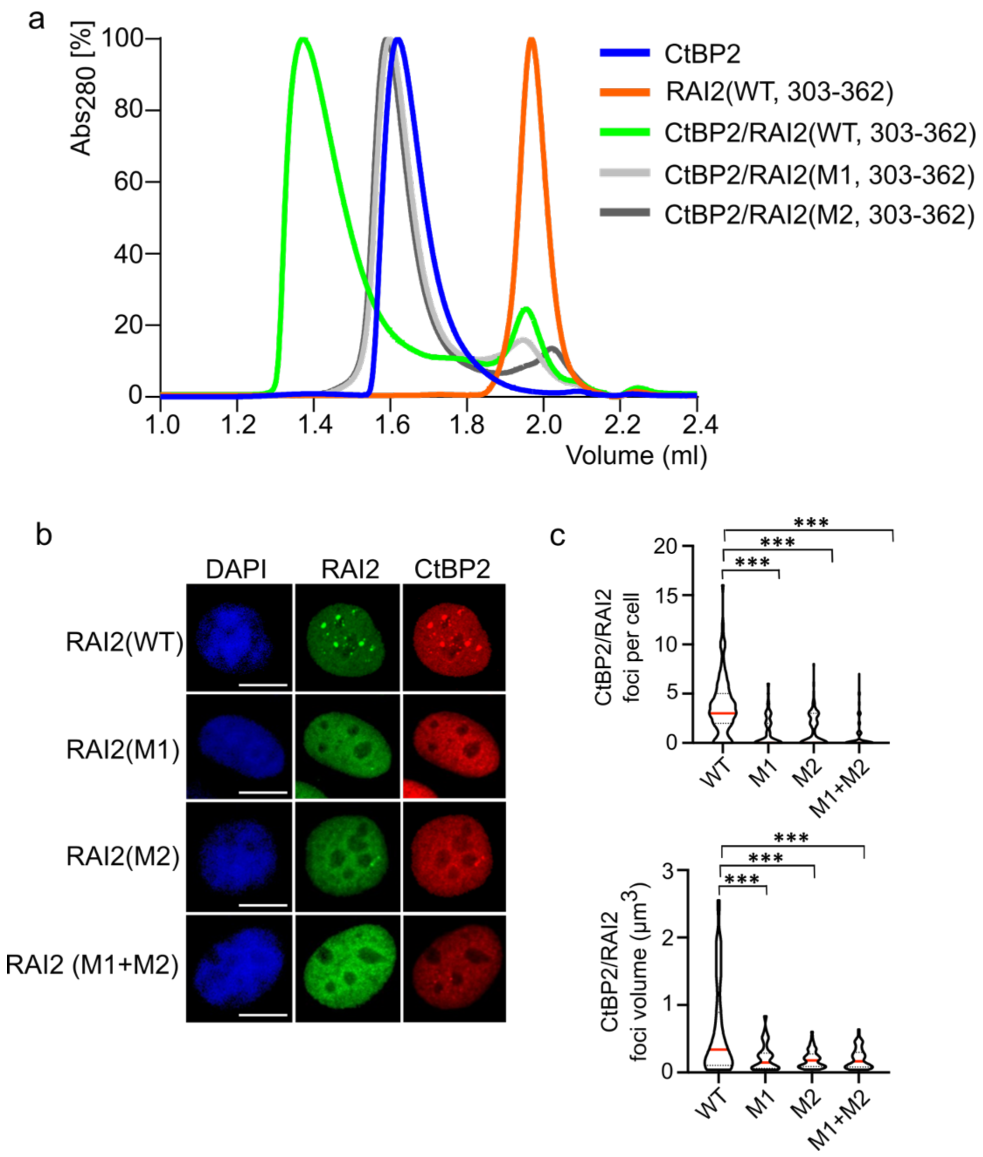

**Supplementary Fig. 7: Characterization of CtBP2/RAI2 complexes**. a. SEC profiles of RAI2(303-362) variants after titration with CtBP2: RAI2(WT), green; RAI2(M1), light grey; RAI2(M2), dark grey. SEC profiles of RAI2 (WT, orange) and CtBP2(blue) alone are also shown. The measured retention volumes are listed in Supplementary Table 2. b. Immunofluorescence co-staining of CtBP2 (red) with HA-tagged RAI2 variants (green) in KPL-1 cell nuclei. Nuclei are stained with DAPI (blue). Scale bars: 10 µm. c. The violin plot depicting the number of CtBP2/RAI2 foci per cell (top) and corresponding volumes (bottom). The medians and quartiles have been shown in red and dotted black lines, respectively. The foci per cell were plotted from 100 cells for each RAI2 construct used. The foci volumes were plotted from n = 62 WT, n = 30, n = 65, n = 68 foci. Unpaired t-test was used to evaluate statistical significance applying p values for a two-sided confidence interval as follows: $p < 0.001$ (***), $p < 0.01$ (**), $p < 0.05$ (*). Definition p values: foci per cell and foci volumes = $p < 0.001$. RAI2(WT) was used as a reference. These data complement our findings on CtBP1/RAI2 assembly (Fig. 2c). Source data are provided as a Source Data file.

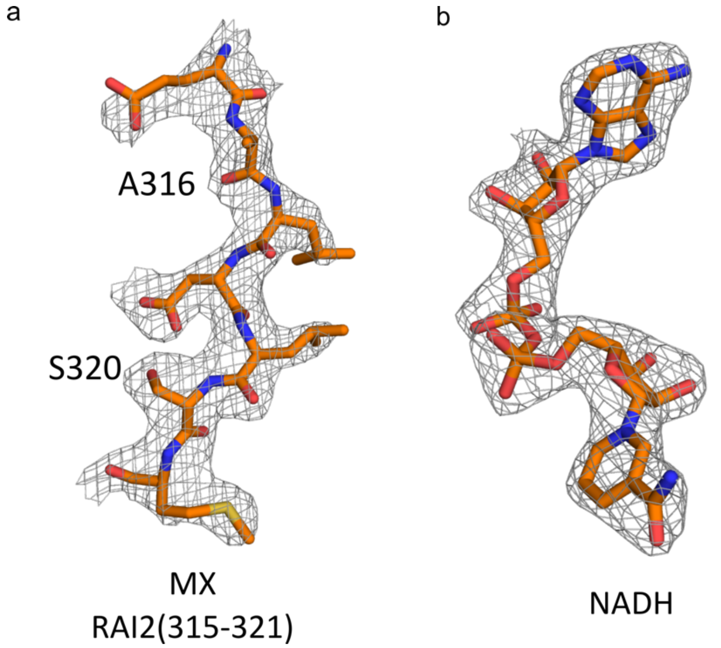

**Supplementary Fig. 8: X-ray crystallography electron density maps of the CtBP2(WT, 31-364)/RAI2(315-321) crystal structure.** a. Single ALDLS motif-containing RAI2 peptide modelled into $2F_o$-$F_c$ electron density at 1σ contour level. b. NADH modelled into omit $2F_o$-$F_c$ electron density at 1σ contour level. Since NCS constraints were used for model refinements the densities are identical in all four CtBP2(WT, 31-364)/RAI2(315-321) complexes.

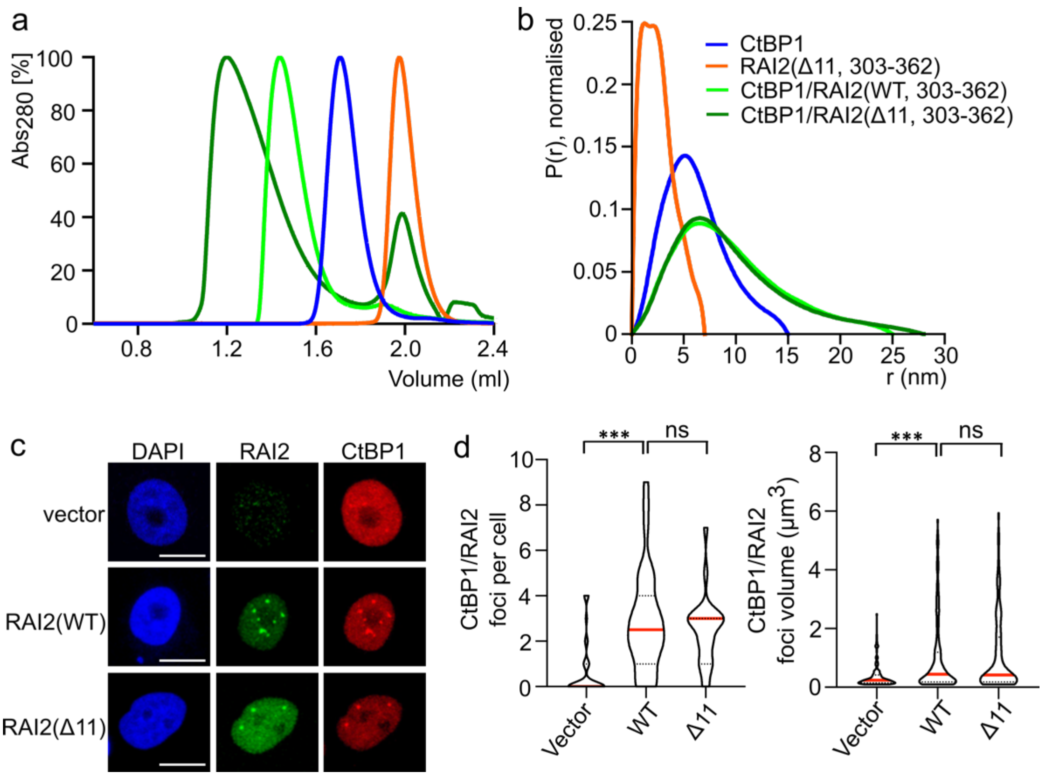

**Supplementary Fig. 9: Biophysical and cellular implications of RAI2(Δ11) on CtBP1 polymerization.** a. SEC profiles after titration of RAI2(303-362) variant titration to CtBP1: CtBP1/RAI2(Δ11), dark green (used for cryo-EM structure determination) CtBP1/RAI2(WT) (green, *cf*. Supplementary Fig. 6d). SEC profiles of RAI2(Δ11, orange) and CtBP1 (blue) alone are also shown. The measured retention volumes are listed in Supplementary Table 2. b. SAXS distance distribution P(r) profiles of CtBP1/RAI2(303-362) variants: CtBP1/RAI2(Δ11), CtBP1/RAI2(WT), CtBP1 alone, RAI2(Δ11) alone, using the same color conventions as in (a). The plots are based on a single experiment and the data have been taken from SASBDB entries SASDQW5, SASDQ76 and SASDQB6. c. Immunofluorescence co-staining of HA-tagged RAI2 variants (Δ11, FL) with CtBP1 in KPL-1 cell nuclei. Co-staining of endogenous RAI2 protein with CtBP1 in vector transduced cells is also shown here for comparison. Nuclei are stained with DAPI (blue). Scale bars: 10 μm. d. Violin plot depicting the number and median of CtBP1/RAI2 foci per cell (left) and their respective volumes (right) of one representative experiment. The medians and quartiles have been shown in red and dotted black lines, respectively. The foci per cell were plotted from n = 37 (vector), n = 34 (WT), n = 23 (Δ11) cells. Foci volumes were plotted from n = 125 (vector), n = 156 (WT), n = 108 (Δ11) foci. Unpaired t-test was used to evaluate statistical significance applying p values for a two-sided confidence interval as follows:

$p < 0.001$ (***), $p < 0.01$ (**), $p < 0.05$ (*). Definition p values: foci per cell: $p < 0.001$ vector, $p = 0.487$ Δ11 and foci volumes: $p < 0.001$ vector, $p = 0.753$ Δ11. CtBP1/RAI2(WT) was used as a reference. Source data are provided as a Source Data file.

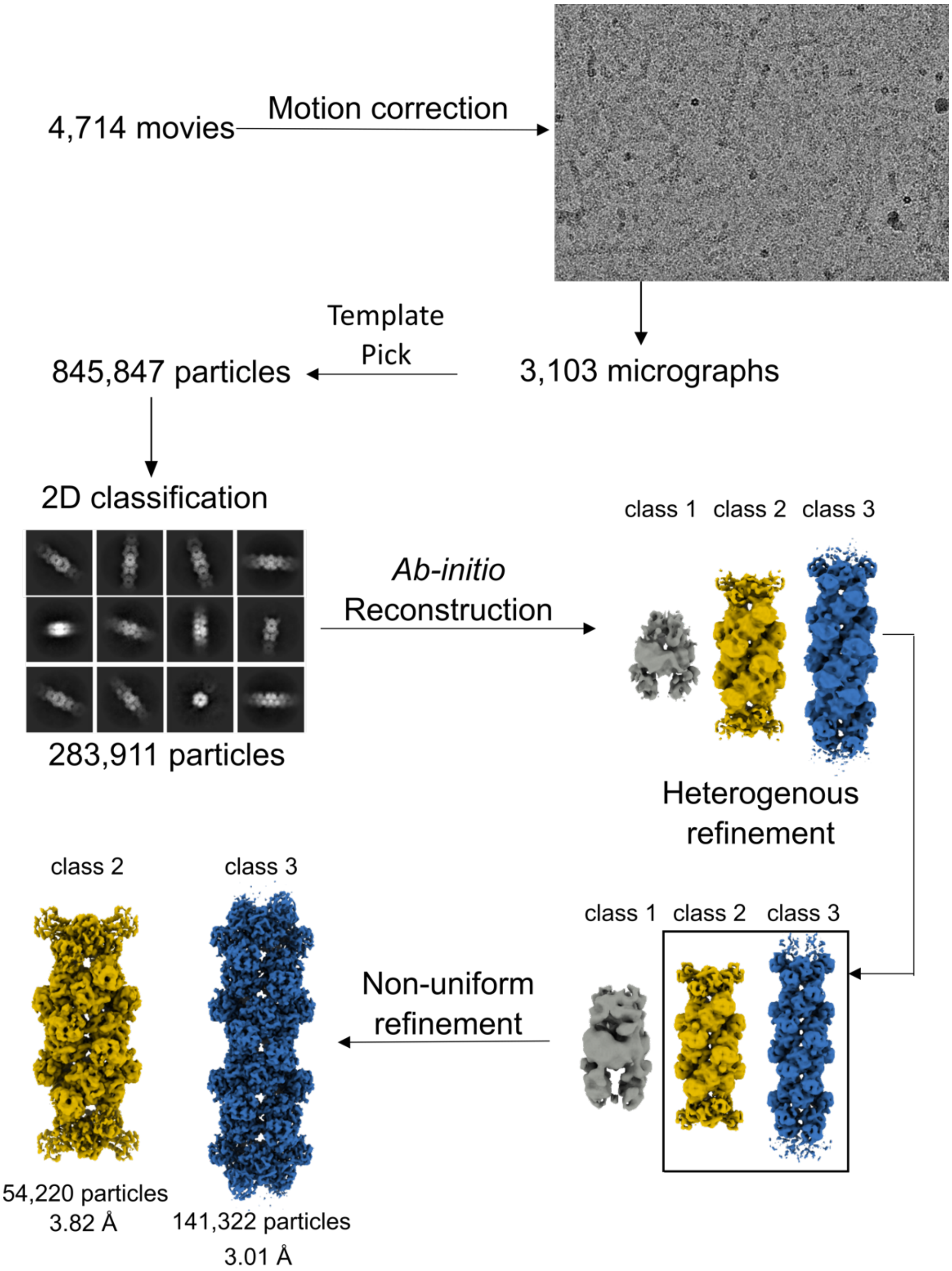

**Supplementary Fig. 10: Cryo-EM data collection and single-particle reconstruction workflow.** Cryo-EM image processing strategy applied to the 4,714 movies in cryoSPARC to generate a D2 symmetry map of the CtBP1/RAI2(Δ11, 303-362) polymeric complex at 3.0 Å resolution. The resulting high-resolution structure is as shown in Fig. 3 and the statistics are listed in Supplementary Table 4.

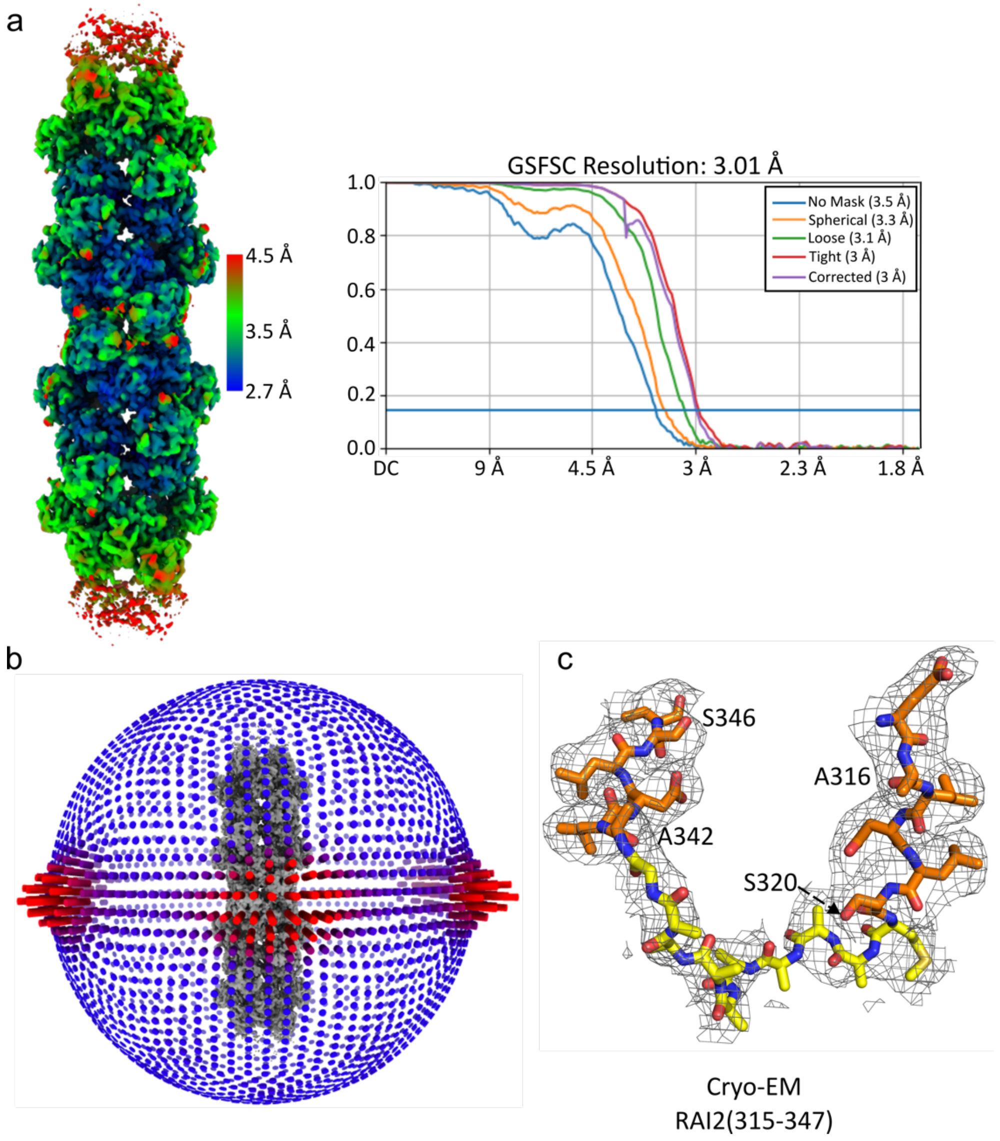

**Supplementary Fig. 11: Quality assessment of the CtBP1(WT, FL)/ RAI2(D11, 303-362) filament cryo-EM structure.** a. The local resolution estimation (left) and the Gold Standard Fourier Shell Correlation (GSFSC) (right) with different masks calculated by cryoSPARC, using a resolution cutoff = 0.143, which is shown as a horizontal line in blue. b. Angular distribution plot for the consensus refinement map. c. Electron density of the central RAI2 peptide segment that includes both ALDLS motifs connecting two adjacent CtBP1 layers (Fig. 3d).

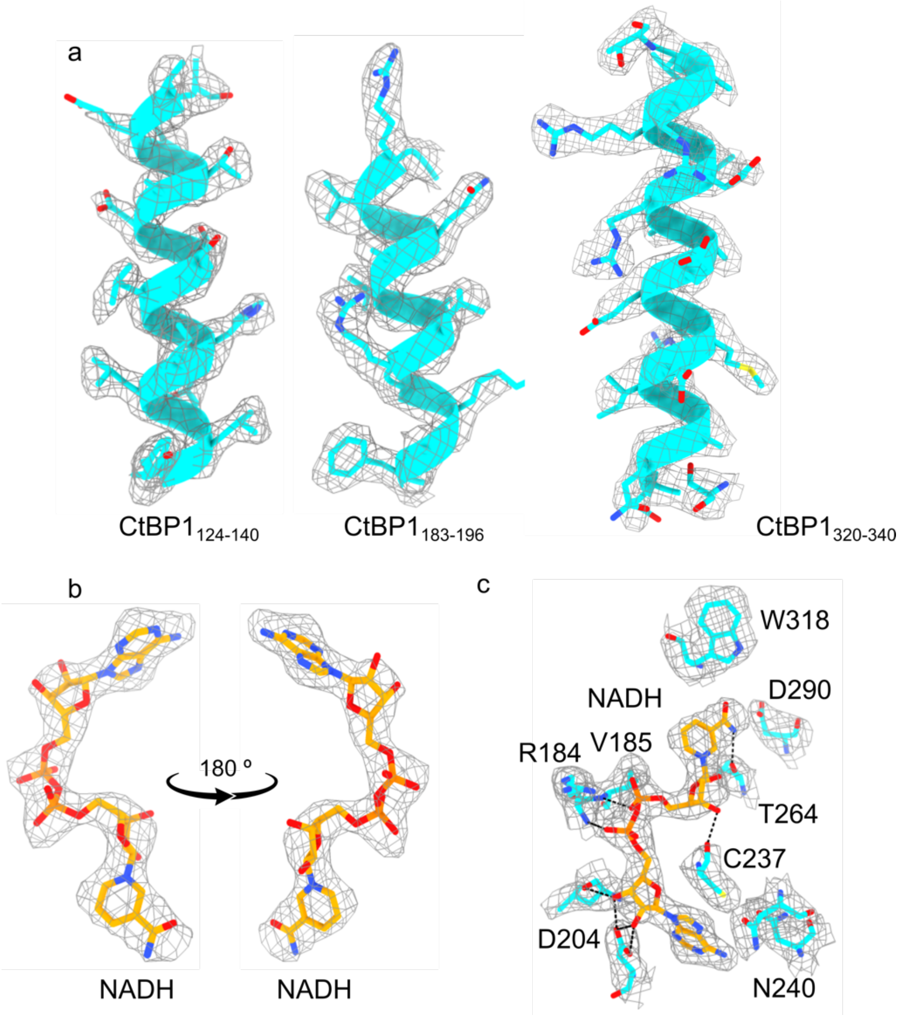

**Supplementary Fig. 12: Representative electron density maps of the CtBP1(WT, FL)/RAI2(D11, 303-362) filament cryo-EM structure.** a. Representative helices of CtBP1 from layer 3 (cf. Fig. 3a). b. NADH shown as stick model at two different viewing angles, rotated by 180° around a vertical axis. c. NADH and interacting CtBP1 residues. Hydrogen bonds are shown as dashed lines. Protein atom colors have been adopted from Fig. 3.

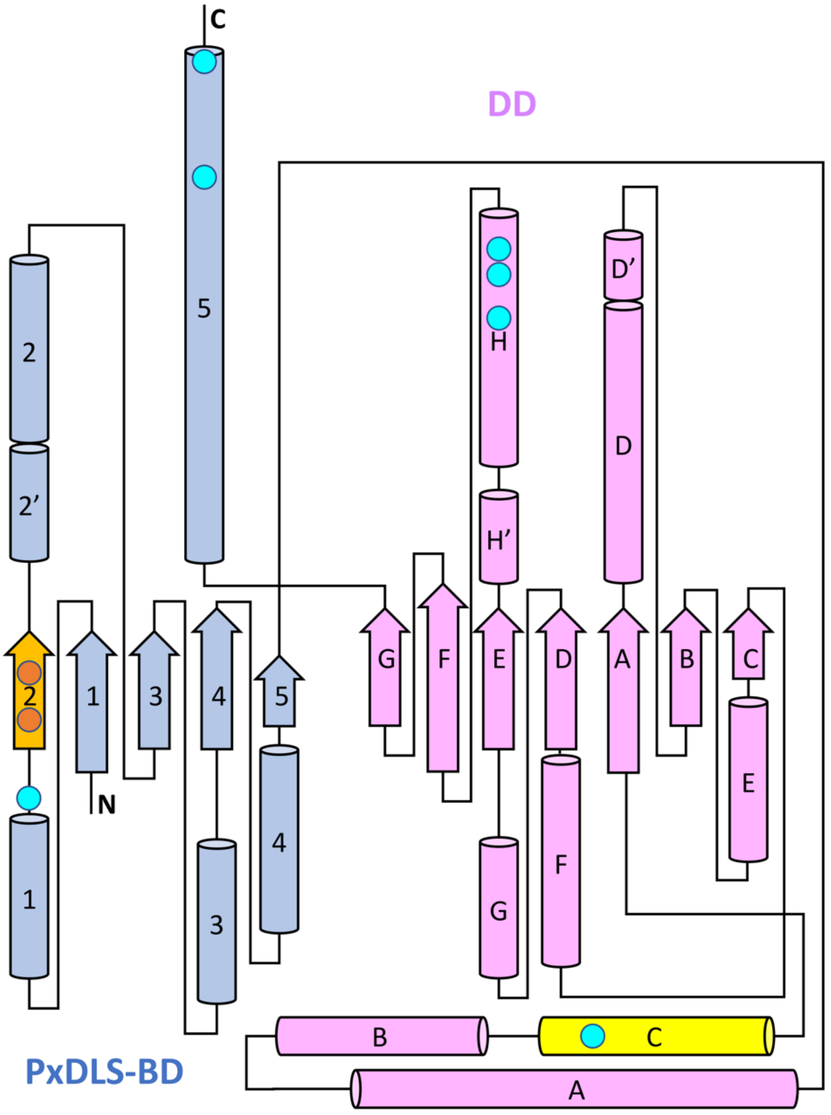

**Supplementary Fig. 13: CtBP subunit topology diagram, covering the PxDLS-BD (blue) and DD (pink), *cf.* Fig.1a.** RAI2-interacting secondary structural elements are in orange (ALDLS motif) and yellow (linker). CtBP residues that contribute to specific interactions (hydrogen bonds) with RAI2 (orange) and fiber CtBP/CtBP layer formation (cyan) are shown by circles. All secondary structure elements are proportional in length. The arrangement of the β-strands corresponds to hydrogen bond pattern of the β-sheet. The RAI2-interacting CtBP β-strand 2 is flanked by the PxDLS-BD β-sheet, to allow the generation of β-sheet like interactions with the two RAI2 ALDLS motifs. Secondary structural element labels are defined as previously [8]. Coordinates of the CtBP1/RAI2(Δ11) filament structure were used for generation of the topology diagram.

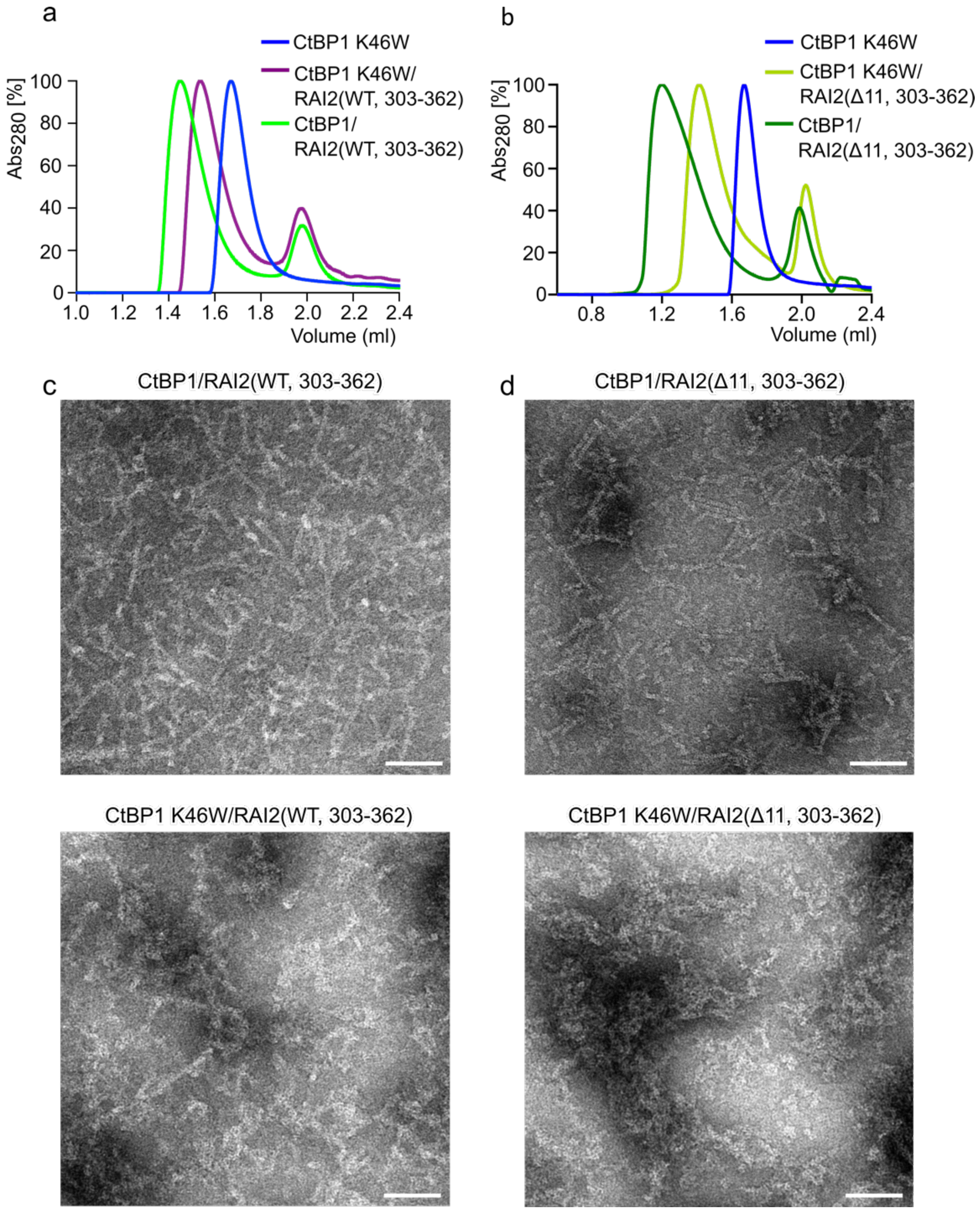

**Supplementary Fig. 14: Effect of the CtBP1 K46W mutation on RAI2/CtBP1 polymerization.** a. SEC profiles after titration of the RAI2(WT, 303-362) (purple) to the CtBP1(K46W) mutant (blue), showing delayed elution volume compared to the WT complex (green, *cf*. Supplementary Fig. 6d). b. SEC profile after titration of the RAI2(Δ11, 303-362) (light green) to CtBP1(K46W) mutant (blue) showing delayed elution retention compared to CtBP1(WT)/RAI2(Δ11, 303-362) complex, *cf.* Supplementary Fig. 9a. The measured retention volumes are listed in Supplementary Table 2. c. Negative staining EM micrographs depicting the CtBP1(WT)/RAI2(WT, 303-

362) complex (top) and CtBP1(K46W)/RAI2 (WT, 303-362) complex (bottom), indicating the loss of the ordered filament due to the mutation. d. Negative stained EM micrographs of the same complexes, using RAI2(Δ11, 303-362) instead of RAI2(WT, 303-362). Scale bars: 100 nm. The micrographs from (c) and (d) are based on a single experiment. Source data are provided as a Source Data file.

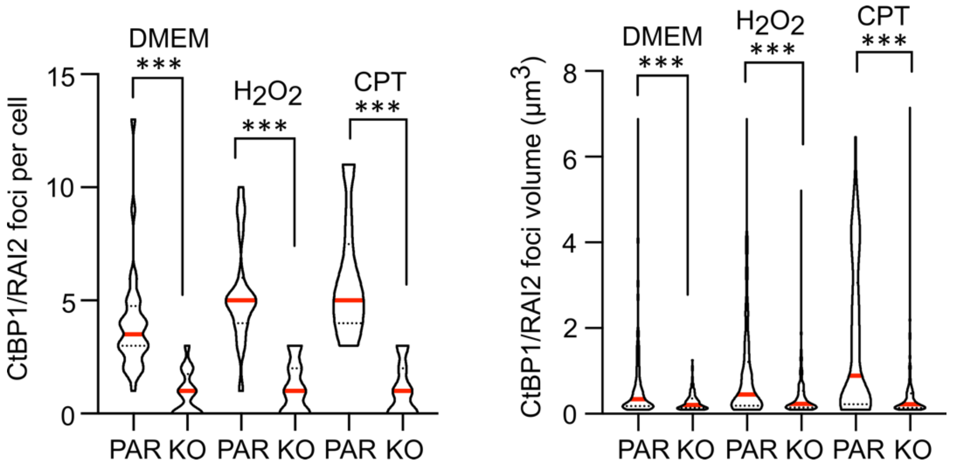

**Supplementary Fig. 15: Effect of genotoxic stress on CtBP1/RAI2 foci in VCaP PC cells.** Violin plot depicting the quantification of CtBP1/RAI2 foci numbers (left) and volumes (right) in PAR and RAI2 KO VCaP cells, untreated (DMEM), treated with $H_2O_2$ and CPT for 48h of one representative experiment (*cf*. Fig. 4). The data are represented as mean of n = 3 independent biological experiments ± SD. The medians and quartiles have been shown in red and dotted black lines, respectively. The foci per cell were plotted from n = 32 (PAR and KO DMEM), n = 19 (PAR $H_2O_2$), n = 18 (KO $H_2O_2$), n = 21 (PAR CPT), n = 18 (KO CPT) cells. Foci volumes were plotted from n = 445 (PAR DMEM), n = 127 (KO DMEM), n = 134 (PAR $H_2O_2$), n = 242 (KO $H_2O_2$), n = 118 (PAR CPT), n = 257 (KO CPT) foci. Unpaired t-test was used to evaluate statistical significance applying p values for a two-sided confidence interval as follows: $p < 0.001$ (***), $p < 0.01$ (**), $p < 0.05$ (*). Definition p values: $p < 0.001$ for all the above-mentioned conditions. The reference data set is the untreated VCaP PAR cell line cultured in DMEM. Source data are provided as a Source Data file.

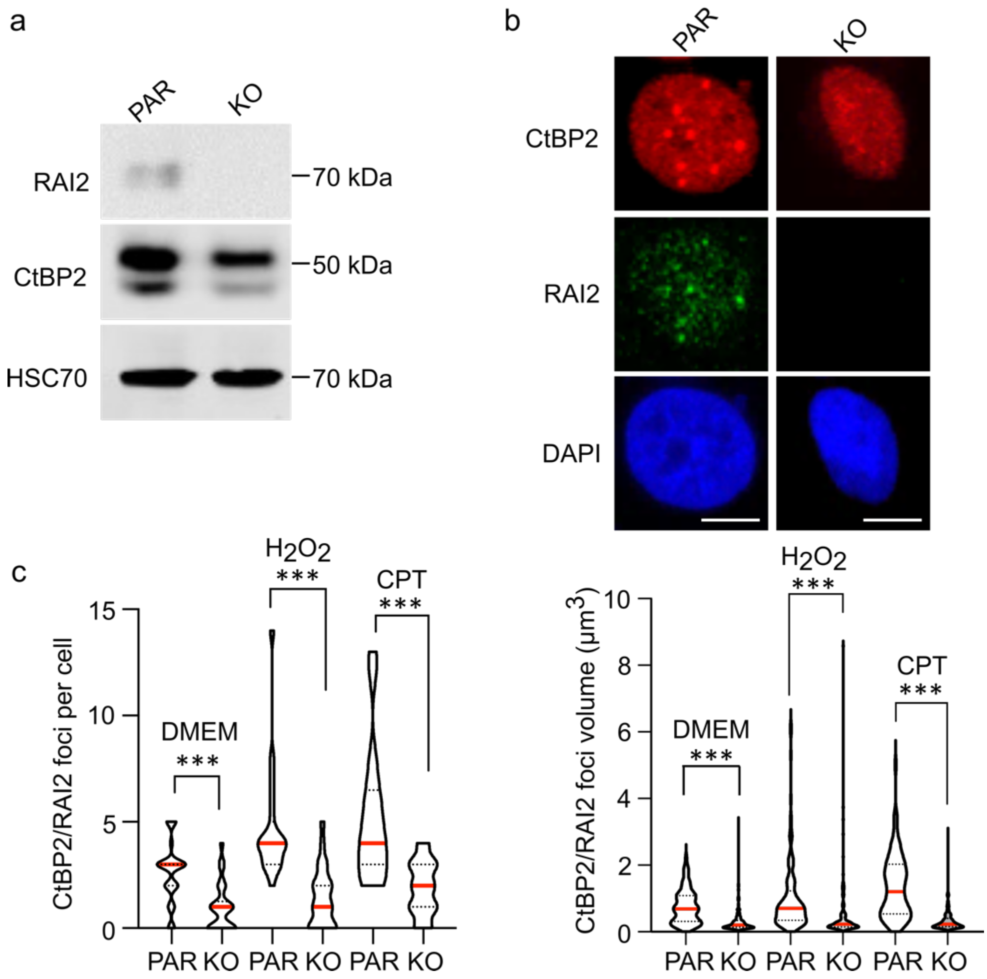

**Supplementary Fig. 16: Effect of genotoxic stress on CtBP2/RAI2 foci in VCaP PC cells.** a. Expression levels of RAI2, CtBP2 and HSC70 (loading control) in PAR and RAI2 KO VCaP PC cells. Biological replicates are provided as a source data file. b. Immunofluorescence staining of CtBP2 (red) and RAI2 (green) in PAR and RAI2 KO VCaP cell nuclei; DAPI (blue) staining was used for localization of nuclei. Scale bars: 10 µm. c. Violin plot depicting the quantification of CtBP2/RAI2 foci numbers (left) and volumes (right) in PAR and RAI2 KO VCaP cells, untreated (DMEM), treated with $H_2O_2$ and CPT for 48h of one representative experiment (*cf.* Fig. 4). The data are represented as mean of n = 3 independent biological experiments ± SD. The medians and quartiles have been shown in red and dotted black lines, respectively. The foci per cell were plotted from n = 30 (PAR DMEM), n = 26 (KO DMEM), n = 19 (PAR $H_2O_2$), n = 26 (KO $H_2O_2$), n = 17 (PAR CPT), n = 20 (KO CPT) cells. The foci volumes were plotted from n = 93 (PAR DMEM), n = 229 (KO DMEM), n = 117 (PAR $H_2O_2$), n = 139 (KO $H_2O_2$), n = 79 (PAR CPT), n = 243 (KO CPT) foci. Unpaired t-test was used to evaluate statistical significance applying p values for a two-sided confidence interval as follows: $p < 0.001$ (***), $p < 0.01$ (**), $p < 0.05$ (*). Definition p values: $p < 0.001$ for all the above-mentioned conditions. The reference data set is the untreated VCaP PAR cell line cultured in DMEM. Source data are provided as a Source Data file.

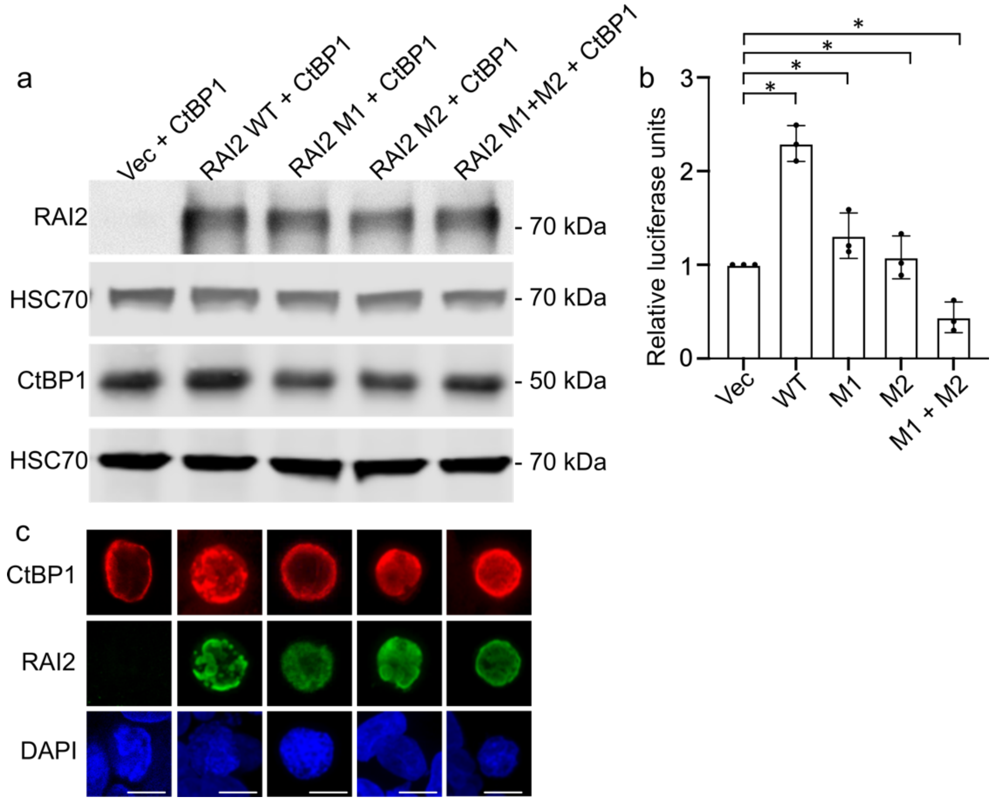

**Supplementary Fig. 17: CDKN1A transactivation assay.** a. Expression levels of RAI2 variants and CtBP1 in transiently transfected HEK 293T cells. Detection of HSC70 serves as loading control. b. Analysis of CDKN1A promoter activation in cells by gene reporter assay in HEK 293T cells transiently transfected with expression plasmids for CtBP1 and RAI2 variants WT, M1, M2 and M1+M2 or empty vector control (Vec). The data are represented as mean of n = 3 independent biological replicates ± SD. Unpaired t-test was used to evaluate statistical significance applying p values for a two-sided confidence interval as follows: $p < 0.001$ (***), $p < 0.01$ (**), $p < 0.05$ (*). Definition p values: $p > 0.001$ WT, $p = 0.005$ M1, $p = 0.002$ M2, $p > 0.001$ M1+M2. Vector control was used as the reference. c. Representative imaging analysis of HEK 293T cells transiently transfected with CtBP1(WT, FL) (red), RAI2(FL) variants (green) and DNA (DAPI, blue). Scale bar: 7.5 µm. Source data are provided as a Source Data file.

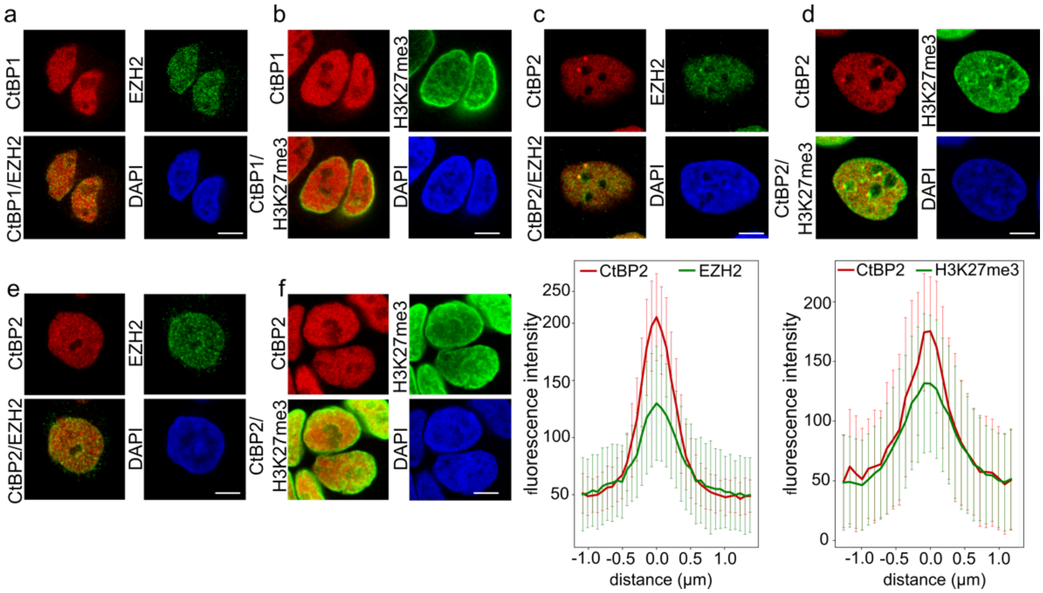

**Supplementary Fig. 18: Colocalization of foci with PRC2 components in VCaP PC cells.** Immunofluorescence staining of VCaP KO cells co-stained for (a) CtBP1 (red), EZH2 (green), CtBP1/EZH2 merge, DNA/DAPI (blue) and (b) CtBP1 (red), H3K27me3 (green), CtBP1/H3K27me3 merge, DNA/DAPI (blue). Immunofluorescence staining of VCaP PAR cells (upper panels) co-stained for (c) CtBP2 (red), EZH2 (green), CtBP2/EZH2 merge, DNA/DAPI (blue) and (d) CtBP2 (red), H3K27me3 (green), CtBP2/H3K27me3 merge, DNA/DAPI (blue). Mean fluorescence intensity profiles ± SD of CtBP2 overlayed with EZH2 or H3K27me3 images (bottom right panels) by line scanning of 100 foci each over 3 µm distance. A total of 100 foci from one representative experiment were analyzed for each condition. Colors are as in the images used for analysis. Immunofluorescence staining of VCaP KO cells co-stained for (e) CtBP2 (red), EZH2 (green), CtBP2/EZH2 merge, DNA/DAPI (blue) and (f) CtBP2 (red), H3K27me3 (green), CtBP2/H3K27me3 merge, DNA/DAPI (blue). Scale bars: 10 µm. Source data are provided as a Source Data file.

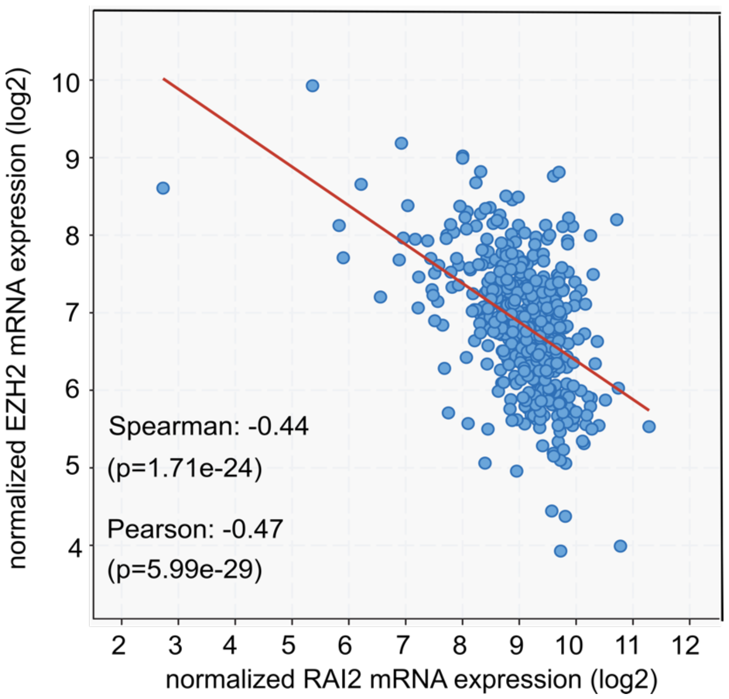

**Supplementary Fig. 19: Correlation and linear regression of RAI2 and EZH2 mRNA expression in 494 primary prostate adenocarcinomas from the TCGA PanCancer Atlas.** mRNA expression is expressed as batch-normalized and log2-transformed value from Illumina HiSeq_RNASeqV2 data. Source data are provided as a Source Data file.

### Supplementary Tables

**Supplementary Table 1: Quantitative binding values of CtBP1/RAI2 variants.**

| Protein complexes | $K_D$(μM) | Student *t*-test P value | Stoichiometry (N, measured) | Stoichiometry (N, interpreted) |
|---|---|---|---|---|
| CtBP1(WT)/ RAI2(WT, 303-362) | 1.1 ± 0.22 | | 0.42 ± 0.04 | 0.5 |
| CtBP1(WT)/ RAI2(WT, 303-465) | 1.5 ± 0.00 | 0.0701 (ns) | 0.32 ± 0.01 | 0.5 |
| CtBP1(WT)/ RAI2(WT, 303-530) | 1.5 ± 0.60 | 0.2960 (ns) | 0.34 ± 0.02 | 0.5 |
| CtBP1(WT)/ RAI2(M1, 303-362) | 5.5 ± 1.00 | 0.0027 (*) | 0.78 ± 0.02 | 1.0 |
| CtBP1(WT)/ RAI2(M2, 303-362) | 3.9 ± 0.46 | 0.004 (*) | 0.84 ± 0.20 | 1.0 |
| CtBP1(WT)/ RAI2(Δ11, 303-362) | 0.8 ± 0.15 | 0.2165 (ns) | 0.41 ± 0.17 | 0.5 |
| CtBP1(K46W)/ RAI2(Δ11, 303-362) | 7.2 ± 0.36 | 0.0021 (*) | 0.29 ± 0.04 | n.d. |
| CtBP1(K46W)/ RAI2(WT, 303-362) | 4.5 ± 0.42 | 0.0032 (*) | 0.26 ± 0.01 | n.d. |

**Legend Supplementary Table 1**: The dissociation constant ($K_D$) and stoichiometry (N) values are represented as a mean of n =3 independent biological experiments + SD. For $K_D$ values, unpaired Student t-test was used to evaluate statistical significance applying p values for a two-tailed confidence interval as follows: $p < 0.001$ (***), $p < 0.01$ (**), $p < 0.05$ (*), “ns” indicates a non-significant p value. Reference data set is CtBP1(WT)/RAI2(WT, 303-362). Source data are provided as a Source Data file.

**Supplementary Table 2: Size exclusion chromatography of CtBP/RAI2 variants.**

| Protein constructs | Elution volume maxima (ml) | Stoichiometry |
|---|---|---|
| CtBP1(WT) | 1.71 | $(CtBP1)_4$ |
| CtBP2(WT) | 1.62 | $(CtBP2)_4$ |
| CtBP1(K46W) | 1.68 | $(CtBP1)_4$ |
| RAI2(WT, 303-362) | 1.98 | $(RAI2)_1$ |
| RAI2(WT, 303-465) | 1.82 | $(RAI2)_1$ |
| RAI2 (Δ11, 303-362) | 2.03 | $(RAI2)_1$ |
| CtBP1(WT)/RAI2(WT, 303-362) | 1.44 | $[(CtBP1)_4RAI2(WT)_2]_x$ |
| CtBP1(WT)/RAI2(M1, 303-362) | 1.69 | $(CtBP1)_4RAI2(M1)_4$ |
| CtBP1(WT)/RAI2(M2, 303-362) | 1.68 | $(CtBP1)_4RAI2(M2)_4$ |
| CtBP1(WT)/RAI2(Δ11, 303-362) | 1.20 | $[(CtBP1)_4RAI2(\Delta11)_2]_x$ |
| CtBP1(K46W) /RAI2(WT, 303-362) | 1.54 | n.d. |
| CtBP1(K46W) /RAI2(Δ11, 303-362) | 1.41 | n.d. |
| CtBP1(WT)/RAI2(WT, 303-465) | 1.36 | $[(CtBP1)_4RAI2(WT)_2]_x$ |
| CtBP1(WT)/RAI2(M1, 303-465) | 1.53 | $(CtBP1)_4RAI2(M1)_4$ |
| CtBP1(WT)/RAI2(M2, 303-465) | 1.50 | $(CtBP1)_4RAI2(M2)_4$ |
| CtBP2(WT)/RAI2(WT, 303-362) | 1.37 | $[(CtBP1)_4RAI2(WT)_2]_x$ |
| CtBP2(WT)/RAI2(M1, 303-362) | 1.60 | $(CtBP1)_4RAI2(M1)_4$ |
| CtBP2(WT)/RAI2(M2, 303-362) | 1.59 | $(CtBP1)_4RAI2(M2)_4$ |

**Legend Supplementary Table 2:** Elution volumes of SEC profile peak maxima of CtBP1, CtBP2, RAI2 and complexes, obtained from a Superose 6 column (see Methods for details) have been depicted. The estimated retention void volume for this column is 0.6 ml. The estimated assembly stoichiometries are listed. For reasons of simplicity, the presence of $NAD^+$/NADH was not included in the estimated stoichiometries. Source data are provided as a Source Data file.

**Supplementary Table 3: X-ray data and structure determination of CtBP2(31-364)/RAI2(M2, 303-465)**

| | CtBP2(WT, 31-364)/ RAI2(M2, 303-465) |
|---|---|
| **Data collection** | |
| Space group | $P6_122$ |
| Cell dimensions | |
| *a*, *b*, *c* (Å) | 126.7, 126.7, 357.6 |
| $\alpha, \beta, \gamma$ (°) | 90, 90, 120 |
| Resolution (Å) | 2.60 |
| $R_{sym}$ or $R_{merge}$ | 0.037 (0.428) |
| $I / \sigma I$ | 12.9 (1.8) |
| Completeness (%) | 99.9 (100.0) |
| Redundancy | 2 |
| **Refinement** | |
| Resolution (Å) | 2.60 |
| No. reflections | 105932 (10449) |
| $R_{work}$ / $R_{free}$ | 0.225/0.287 |
| No. atoms | |
| Protein | 10387 |
| Ligand/ion | 183 |
| Water | 206 |
| *B*-factors | |
| Protein | 47.9 |
| Ligand/ion | 37.9 |
| Water | 52.8 |
| R.m.s. deviations | |
| Bond lengths (Å) | 0.014 |
| Bond angles (°) | 2.01 |

* Values in parentheses are for highest-resolution shell.

**Supplementary Table 4: Cryogenic-EM data collection, refinement, and validation statistics of CtBP1(WT)/RAI2(Δ11, 303-362) structure**

| | CtBP1(WT)/RAI2(Δ11, 303-362) (EMDB-15603) (PDB 8ARI) |
|---|---|
| **Data collection and processing** | |
| Magnification | 105,000 |
| Voltage (kV) | 300 |
| Electron exposure (e–/Å$^2$) | 42 |
| Defocus range (μm) | 0.75-2.75 |
| Pixel size (Å) | 0.87 |
| Symmetry imposed | D2 |
| Initial particle images (no.) | 845847 |
| Final particle images (no.) | 189823 |
| Map resolution (Å) | 3.01 |
| FSC threshold | 0.143 |
| Map resolution range (Å) | 2.7-4.5 |
| **Refinement** | |
| Initial model used (PDB code) | 6CDF |
| Model resolution (Å) | |
| FSC threshold | |
| Model resolution range (Å) | |
| Map sharpening *B* factor (Å$^2$) | -100 |
| Model composition | |
| Non-hydrogen atoms | 63832 |
| Protein residues | 8160 |
| Ligands | 24 |
| *B* factors (Å$^2$) | |
| Protein | 71.8 |
| Ligand | 60.5 |
| R.m.s. deviations | |
| Bond lengths (Å) | 0.002 |
| Bond angles (°) | 0.439 |
| Validation | |
| MolProbity score | 1.26 |
| Clashscore | 2.37 |
| Poor rotamers (%) | 0.38 |
| Ramachandran plot | |
| Favored (%) | 96.35 |
| Allowed (%) | 3.52 |
| Disallowed (%) | 0.12 |
