## Supplementary Data 8 for "Master corepressor inactivation through multivalent SLiM-induced polymerization mediated by the oncogene suppressor RAI2"

sp|P0558_RAI2_full_length|P0558_RAI2_full_length − protein_id: P0558_RAI2_full_length − sequence coverage: 69.5 % − total score: 9302

180222_band02_R1

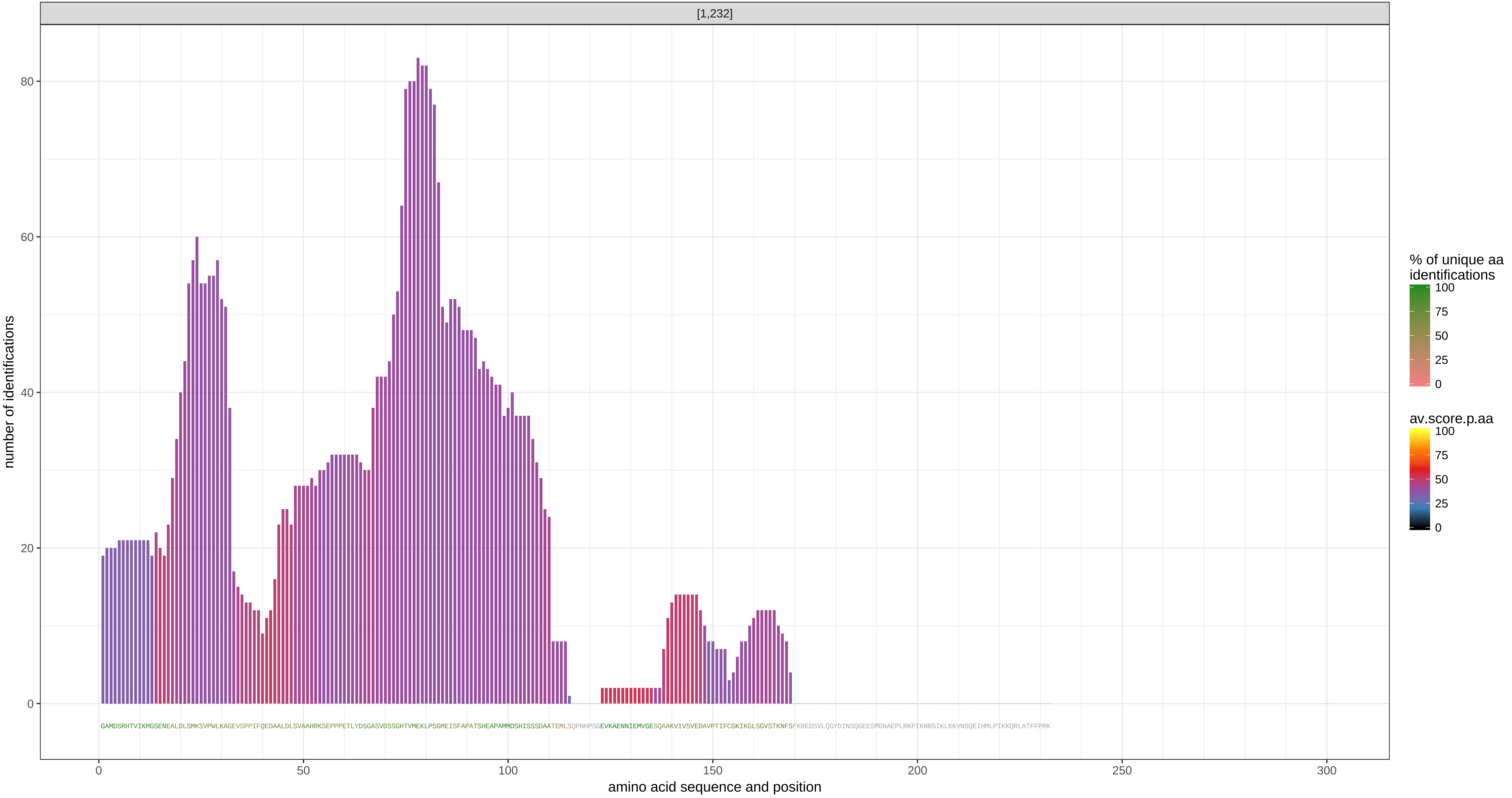

KRT9 − protein_id: ###CONTAMINANT###P35527 − sequence coverage: 22.1 % − total score: 5792

180222_band02_R1

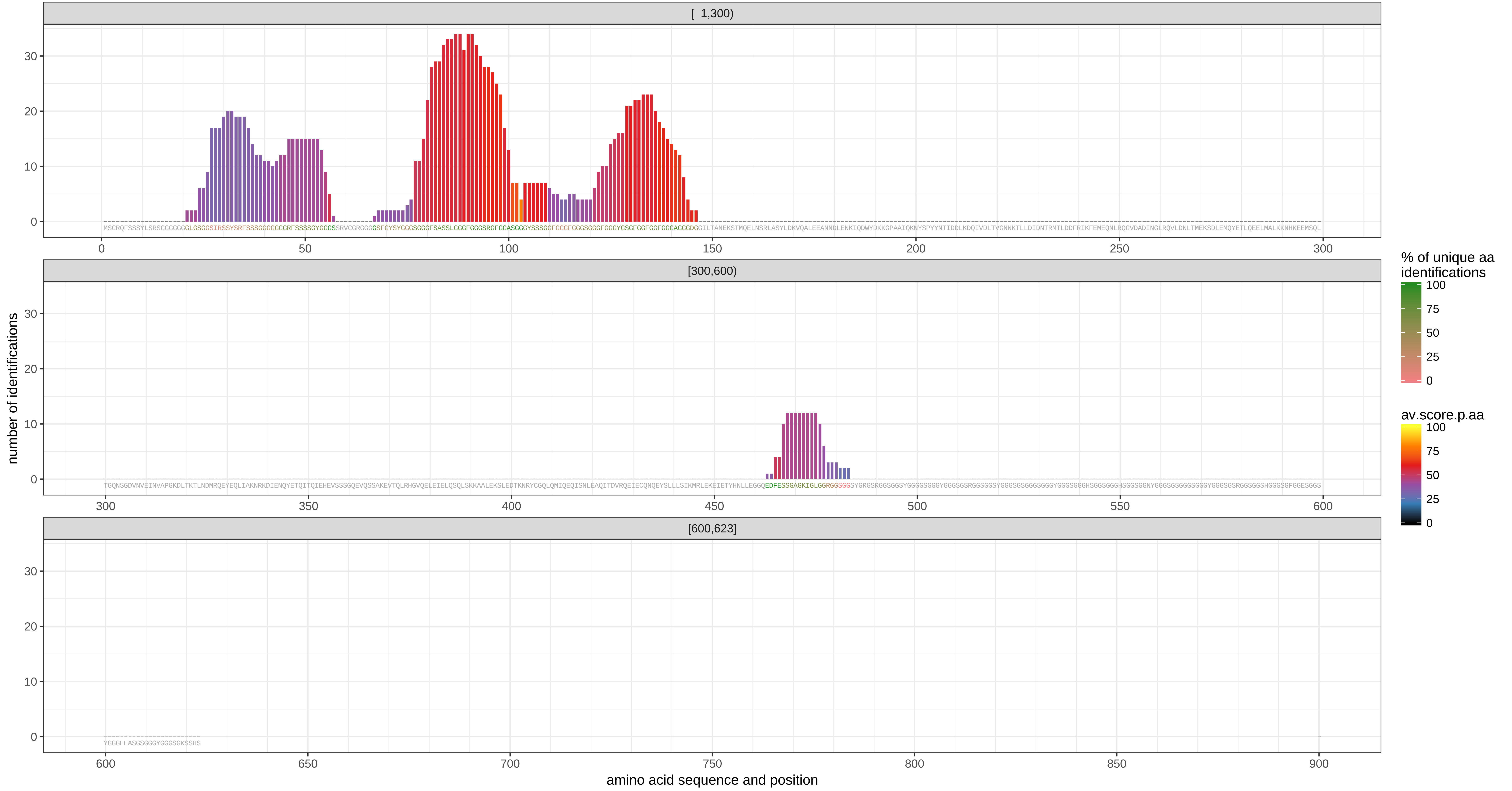

KRT1 − protein_id: ###CONTAMINANT###P04264 − sequence coverage: 24.2 % − total score: 4534

180222_band02_R1

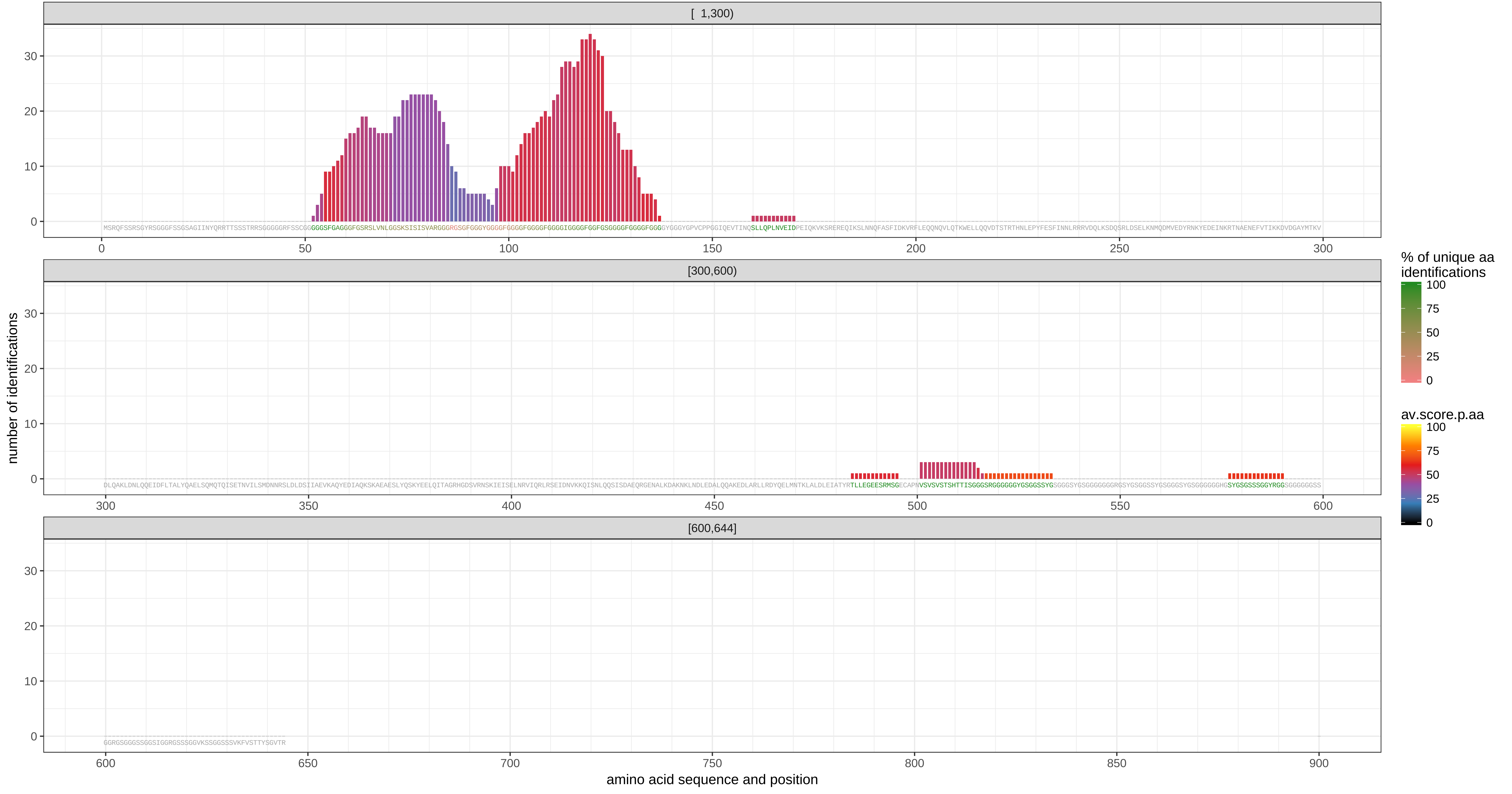

KRT2 − protein_id: ###CONTAMINANT###P35908 − sequence coverage: 13.6 % − total score: 1643

180222_band02_R1

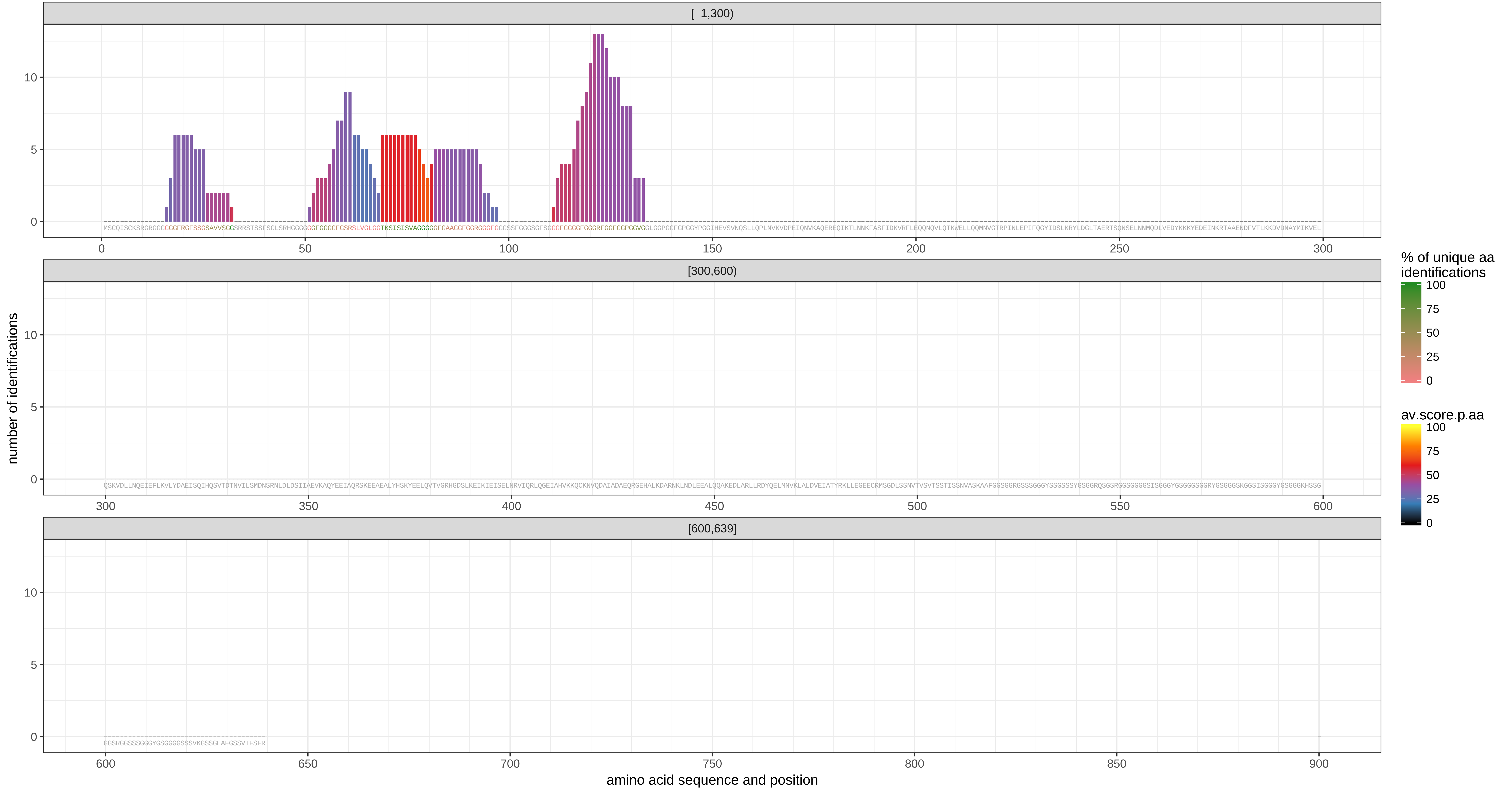

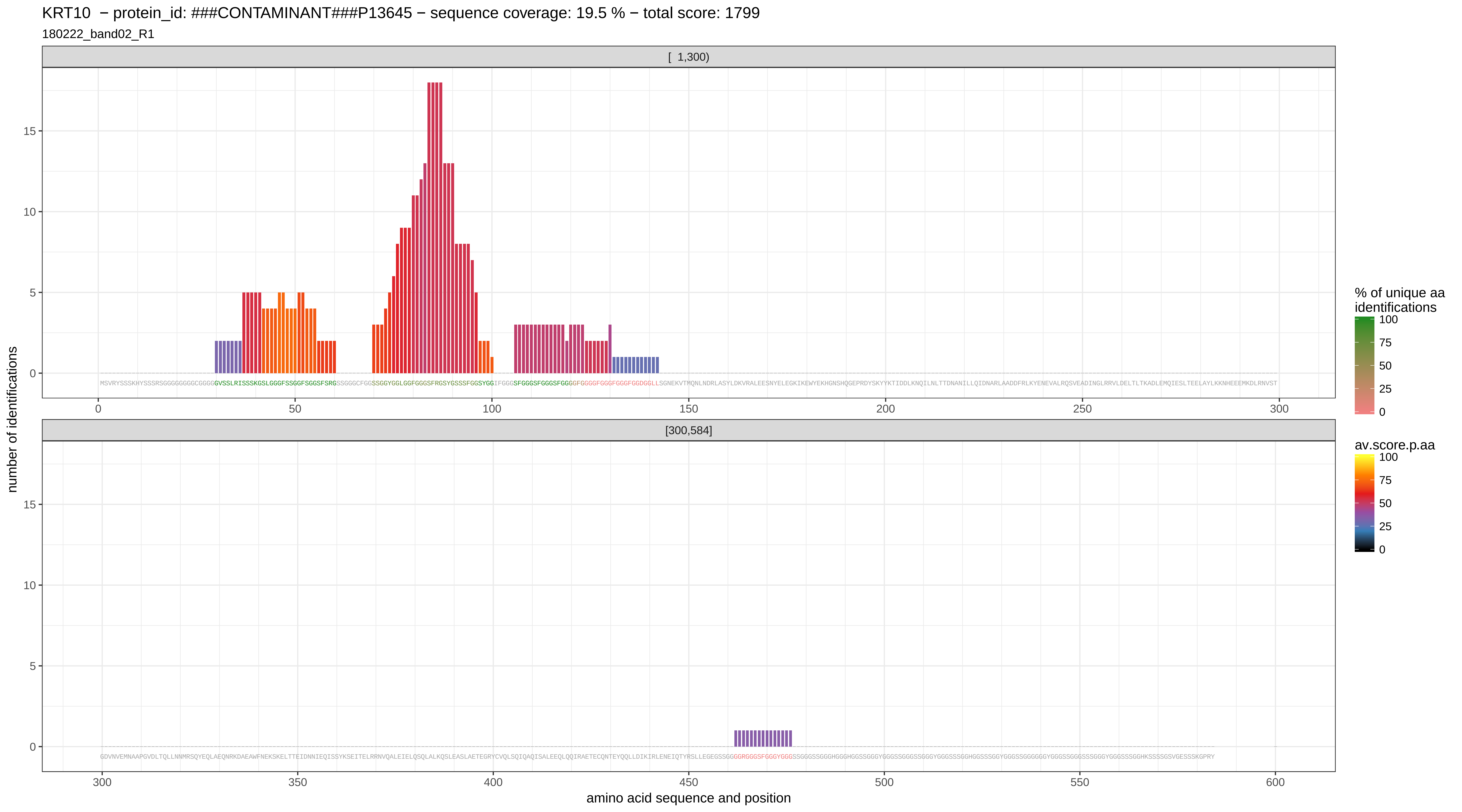
KRT10 − protein_id: ###CONTAMINANT###P13645 − sequence coverage: 19.5 % − total score: 1799
180222_band02_R1
[ 1,300)
[300,584]
number of identifications
amino acid sequence and position
% of unique aa identifications
av.score.p.aa

KRT13 − protein_id: ###CONTAMINANT###P13646|###REV######CONTAMINANT###P13646 − isoform: ###CONTAMINANT###P13646 − sequence coverage: 3.3 % − total score: 200

180222_band02_R1

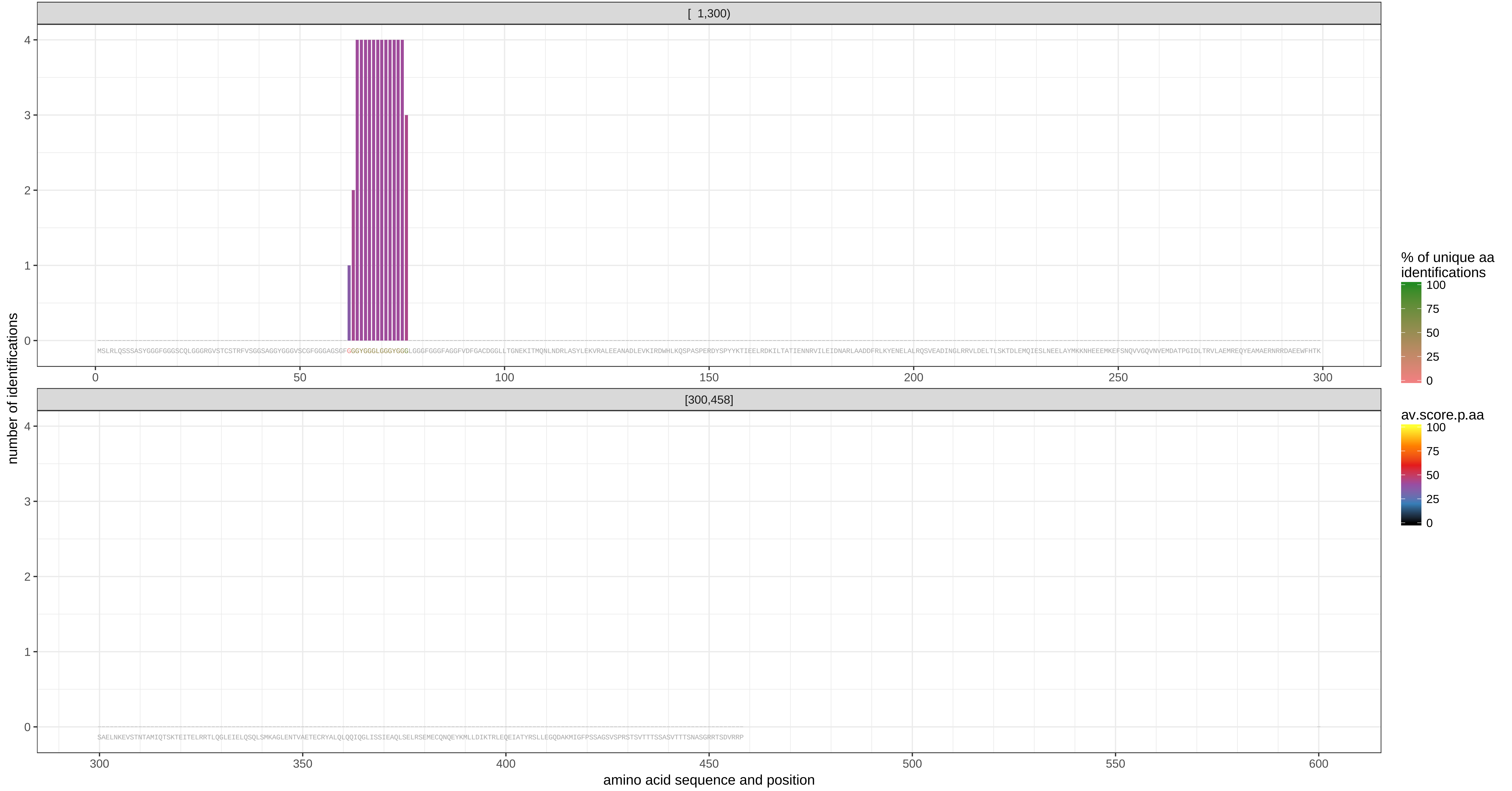

− protein_id: ###CONTAMINANT###P13646|###REV######CONTAMINANT###P13646 − isoform: ###REV######CONTAMINANT###P13646 − sequence coverage: 6.1 % − total score: 200

180222_band02_R1

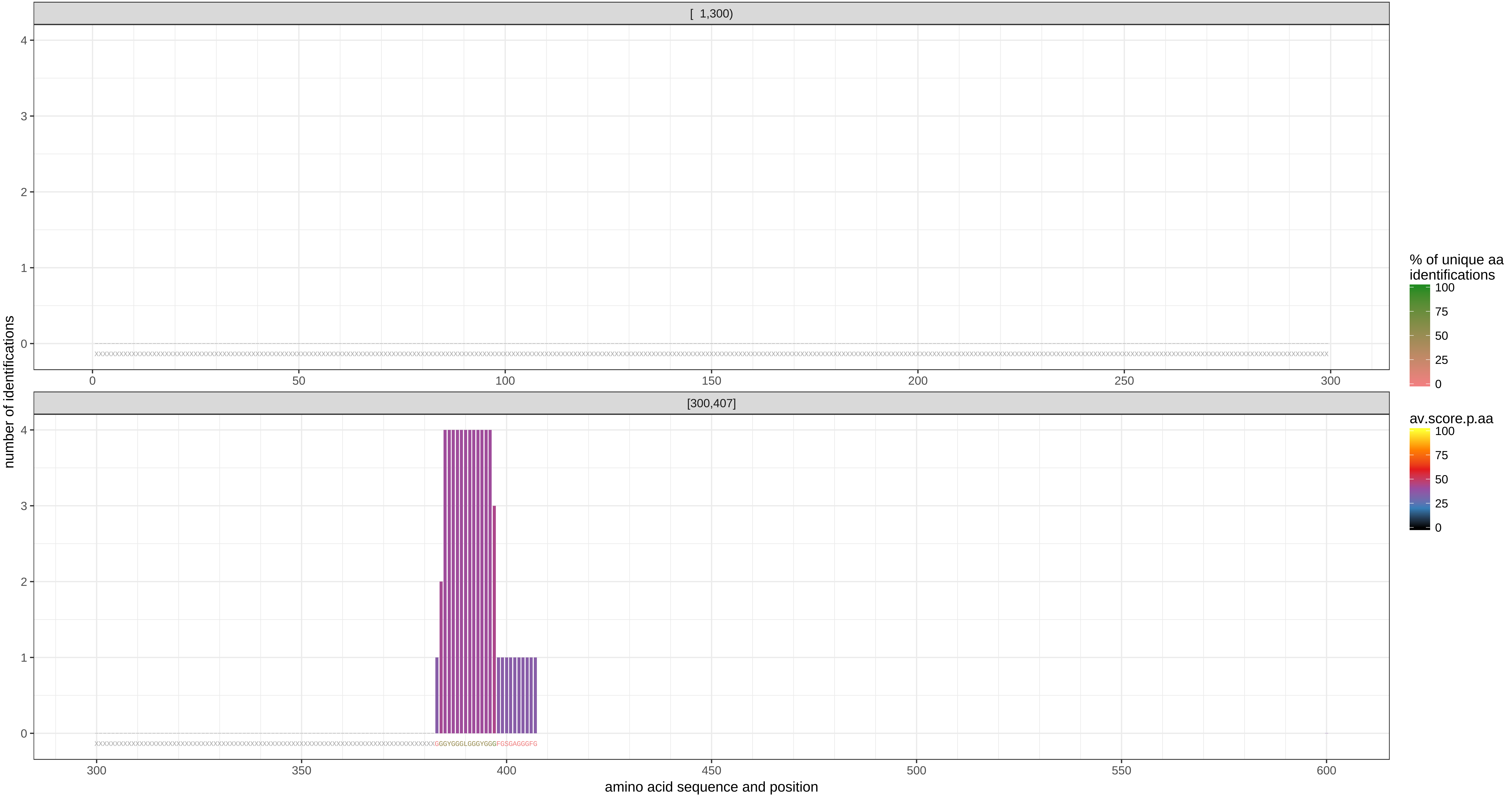

− protein_id: ###REV######CONTAMINANT###P13645 − sequence coverage: 2.8 % − total score: 220

180222_band02_R1

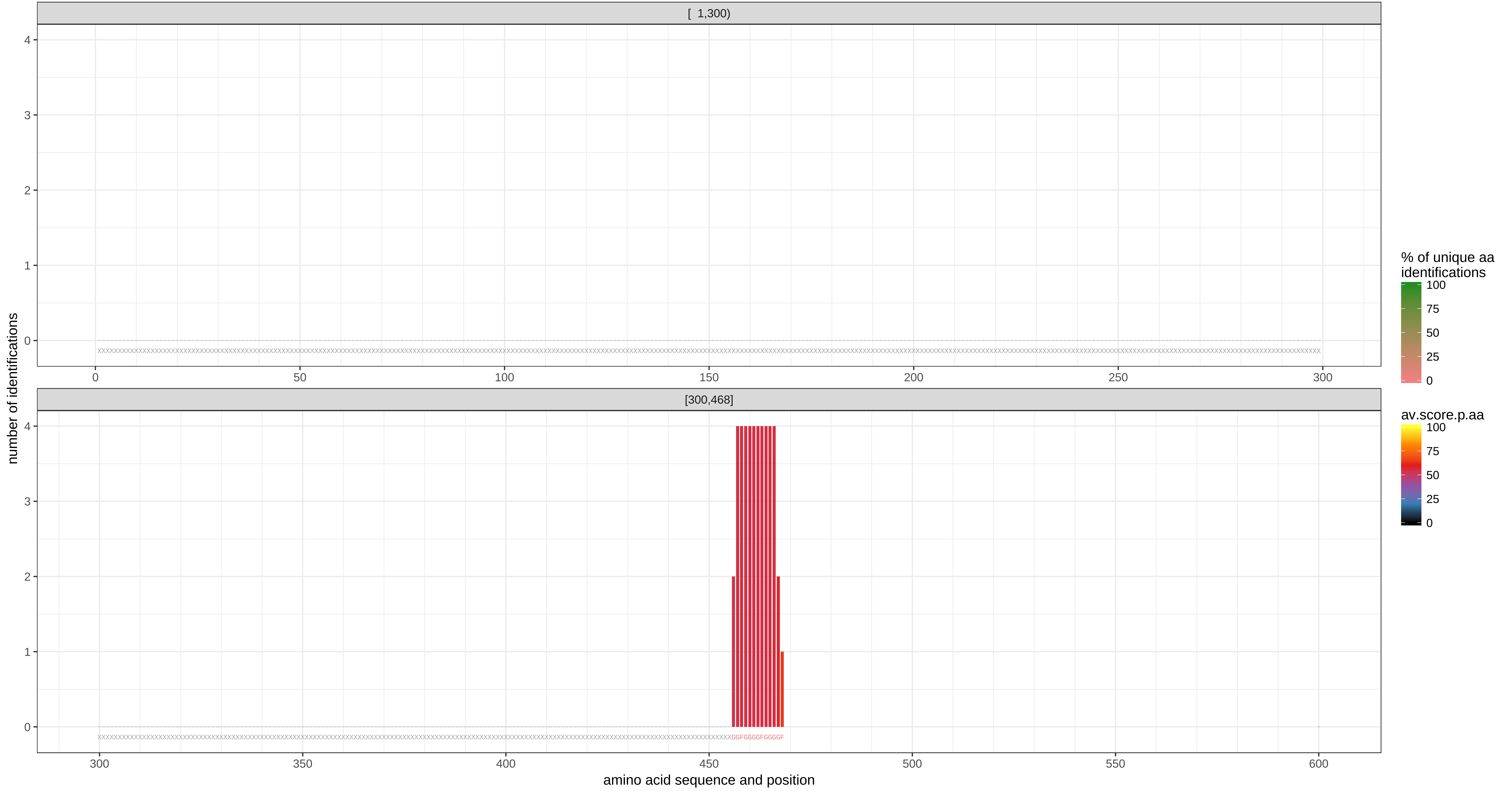

− protein_id: ###REV######CONTAMINANT###P35908 − sequence coverage: 5.7 % − total score: 183

180222_band02_R1

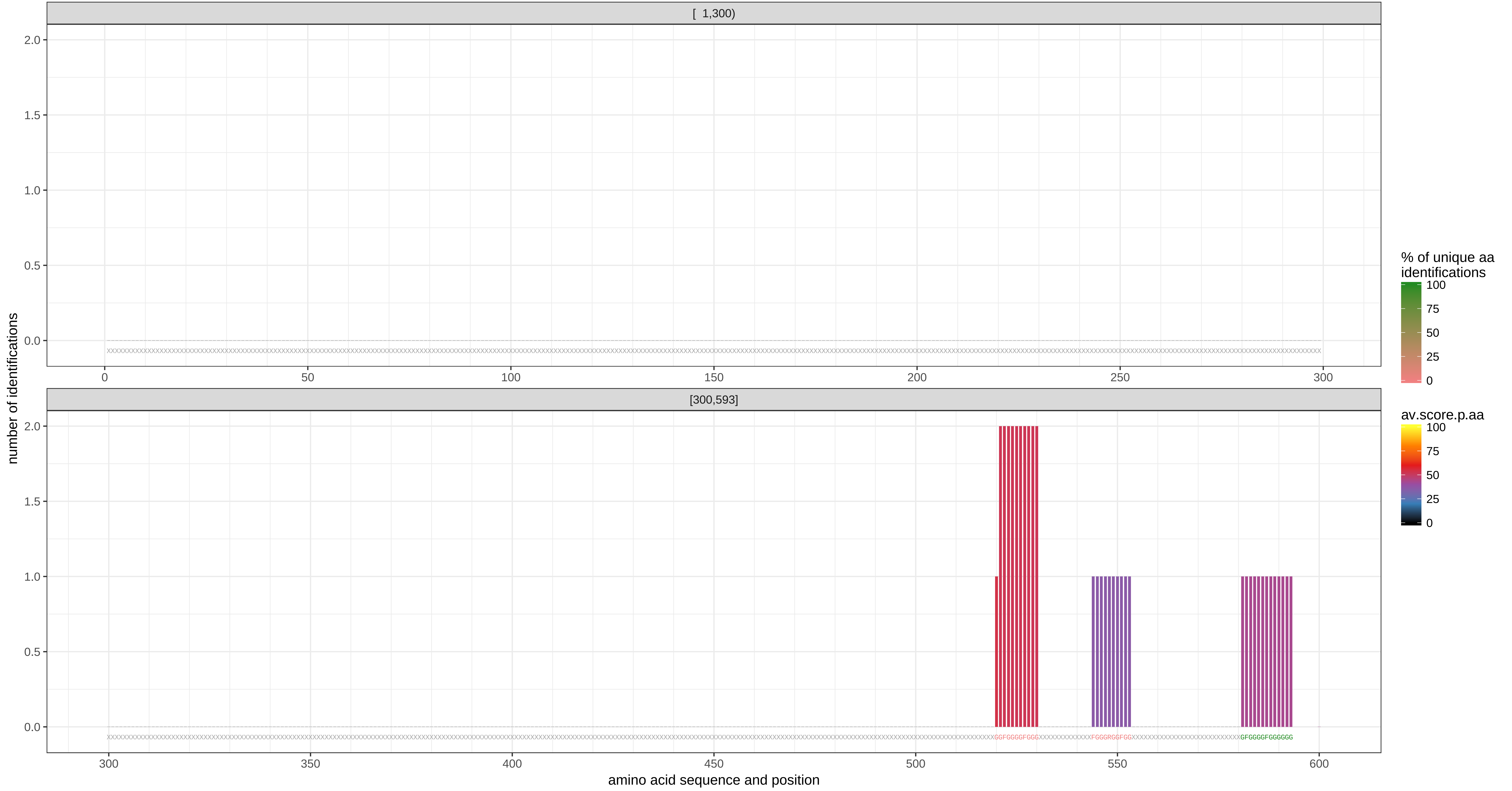
