## Supplementary material for "Master corepressor inactivation through multivalent SLiM-induced polymerization mediated by the oncogene suppressor RAI2": Description of Supplementary Data

### Description of Supplementary Data Files

**Supplementary Data 1: SAXS data collection parameters and analysis of RAI2(303-465) constructs.**

**Supplementary Data 2: SAXS data collection parameters and analysis of RAI2(303-362) constructs.**

**Supplementary Data 3: Nuclear Magnetic Resonance spectroscopy of RAI2(303-362) proteins.** $^{1}H$ and $^{15}N$ chemical shift assignments of RAI2 (WT, 303-362), RAI2(M1, 303-362) and RAI2(M2, 303-362) proteins in ppm have been shown. The data for the two ALDLS motifs are in orange while the mutated residues in M1 and M2 are in red. Residues with overlapping signals are marked with an asterisk.

**Supplementary Data 4: NMR analysis of CtBP1(WT, 28-353) titration to WT, M2 and M1 constructs of RAI2(303-362).** Normalized intensity ($I$) ratios calculated by $I_{complex}/I_{free}$ for each residue of RAI2 WT, M1 and M2 variants are shown. Normalization was carried out by setting the ratio of the unperturbed residue D362 at 1.000. Low intensity ratios indicate perturbation of RAI2 residues in the presence of CtBP. The ratios for WT RAI2 are shown as histogram in **Fig. 1h** top panel. The M1 and M2 ratios were used to generate the histograms in the middle and bottom panels of **Fig. 1h**. The data for the two ALDLS motifs are in orange while the mutated residues in the RAI2 M1 and M2 variants are in red.

**Supplementary Data 5: SAXS Data collection parameters and analysis of CtBP1/RAI2(303-465) complexes.**

**Supplementary Data 6: SAXS Data collection parameters and analysis of CtBP1/RAI2(303-362) complexes.**

**Supplementary Data 7: Gene expression analysis of CTCs isolated from prostate cancer patient blood samples.** Gene expression values of the androgen receptor (*AR*), keratin 19 (*KRT19*) and *RAI2* are expressed as normalized quantification cycles (Cq). Cq values are based on a single experiment using preamplified cDNA from AdnaTest enriched blood samples. Samples were categorized into HSPC, CRPC, AVPC and NEPC groups. Samples with gene expression of *AR* and/or *KRT19* above the defined threshold (see Materials and Methods) were categorized as CTC-positive and are marked with a + sign.

**Supplementary Data 8: Mass Spectrometry analysis of RAI2(WT, 303-530).**
